# Impact of early-life respiratory syncytial virus infection on cell type-specific airway DNA methylation

**DOI:** 10.1101/2024.09.29.615688

**Authors:** Chris McKennan, Dawn C. Newcomb, Sergejs Berdnikovs, Tebeb Gebretsadik, Siyuan Ma, Leonard B. Bacharier, Max A. Seibold, Larry J. Anderson, James E. Gern, Carole Ober, Tina Hartert

**Affiliations:** Department of Statistics, University of Pittsburgh, Pittsburgh, Pennsylvania, USA; Department of Medicine, Vanderbilt University Medical Center, Nashville, Tennessee, USA; Department of Medicine, Northwestern Feinberg School of Medicine, Chicago, Illinois, USA; Department of Biostatistics, Vanderbilt University Medical Center, Nashville, Tennessee, USA; Monroe Carell Jr. Children’s Hospital at Vanderbilt, Division of Pediatric Allergy, Immunology, and Pulmonary Medicine, Nashville, Tennessee, USA; Center for Genes, Environment, and Health at National Jewish Health, Denver, Colorado, USA; Department of Pediatrics, Emory University School of Medicine and Children’s Healthcare of Atlanta, Georgia, USA; Department of Pediatrics, University of Wisconsin School of Medicine and Public Health-Madison, Madison Wisconsin, USA; Department of Human Genetics, University of Chicago, Chicago, Illinois, USA; Department of Pediatrics, Vanderbilt University Medical Center, Nashville, Tennessee, USA

**Keywords:** Respiratory syncytial viruses, DNA methylation, childhood wheeze, childhood asthma

## Abstract

**Rationale:** Infection with respiratory syncytial virus (RSV) in early-life is associated with subsequent childhood respiratory disorders, but the mechanism for this association is unknown. We hypothesized that RSV-mediated alteration in airway DNA methylation (DNAm) may play a role.

**Objectives:** Investigate the impact of early-life RSV infection on airway DNAm.

**Methods:** Our study population consisted of children from the INSPIRE population-based birth cohort. Early-life infection with RSV was defined as infection not requiring clinical illness before age one year, ascertained through active and passive surveillance. Methylation was measured in DNA from nasal airway epithelial cells (NAECs) at ages two years (n=88) and six years (n=539). Bulk gene expression in NAECs was available in a subset of participants at age two years (n=54). *In silico* cell type deconvolution was used to infer cell type-specific associations in suprabasal, ciliated, and “other” cell types.

**Measurements and Main Results:** We identified 164 CpGs in ciliated cells and seven CpGs in suprabasal cells whose cell type-specific DNAm at age two years was associated with early-life RSV infection. The seven associations in suprabasal cells were still present at age six years and the directions of association replicated in *in vitro* infection of epithelial cells with RSV in air liquid interface cultures. DNAm levels at four of the seven CpGs identified in suprabasal cells correlated with the expression of their nearest genes in suprabasal cells, which included genes previously implicated in asthma-related diseases. The DNAm levels of these four CpGs and the expression of their nearest genes in suprabasal cells at age two years were also associated with recurrent wheezing at age four years.

**Conclusions:** We demonstrate epigenetic changes in airway progenitor cells of children who have had RSV infection in the first year of life are conserved over time, associated with subsequent wheeze, have functional relevance, and replicate in *in vitro* infection in ALI cultures. Our study is the first to show that early-life RSV infection, as opposed to severe illness, alters DNAm levels at functionally and clinically relevant loci.

## Introduction

Respiratory syncytial virus (RSV) is a ubiquitous infant and childhood respiratory virus, with roughly half of all children infected by one year of age [1]. While the association between RSV bronchiolitis requiring hospitalization and later childhood asthma is well established, such severe illnesses account for only about 2% of all RSV infections. Further, this association between severe RSV illness and asthma is likely confounded by host genetics [2, 3]. As such, assessing the impact of RSV on childhood respiratory disorders requires studying general RSV infections rather than severe infections. Recently, we showed that the approximately 50% of infants infected with RSV prior to one year, assessed by qPCR or serology, were significantly more likely to develop wheeze or asthma by age five years [1]. While this and other work suggest a causal role for RSV infection in childhood wheeze and asthma, the mechanism(s) is unknown.

DNA methylation (DNAm) is an epigenetic modification that may be involved in the etiology of respiratory diseases. DNAm changes have been associated with exposure histories, including viral infections [4, 5, 6, 7]. We therefore hypothesized that one mechanism through which infant RSV infection may contribute to later childhood respiratory disease is through altering host DNAm patterns in the airway, particularly during early infancy, which represents a crucial period of immune and lung development. This study aims to determine if epigenetic signatures in specific airway epithelial cell subsets differ between children who were and were not infected with RSV before age one year, and whether these changes are associated with subsequent risk for childhood respiratory diseases.

## Methods

### Study participants

The sub-group for this study is an *a priori* designed nested cohort of 100 participants selected for follow-up using a random number generator from 4 pre-defined groups with and without wheezing, and RSV infected and not infected during the first year of life. Wheezing and RSV infection were defined at age one year. The children in this nested cohort completed additional 2-3-year and 6-7 year in-person study visits that included nasal brush sampling used for the DNAm array, nasal airway epithelial cell (NAEC) collection and culture and RNA-seq. In addition, we used nasal airway epithelial cell DNAm array from all available 6-year old children who similarly had known RSV infection status during infancy to assess durability of the DNA methylation signatures identified at age 2-3 years. The clinical outcome of wheeze at four years was defined as two or more wheezing episodes in the past 12 months. The Institutional Review Board of Vanderbilt University approved this study and one parent of each child provided informed consent for their participation.

### NAEC collection for DNA methylation and airway epithelial culture

NAECs were collected from INPSIRE children by trained study nurses at 2-3 and/or 6-7 years of age by brushing the inferior turbinate of the nasal passages with a soft flocked cotton swab (Copan, Murrieta, CA, USA). NAECs brush samples collected at age 2-3 years were cultured in PneumaCult ALI Medium for 3 weeks. Once NAECs were fully differentiated, each of the donor cells were split into 4 groups and infected on the apical surface (top chamber) with 50uL of RSV 2014-1 01/2-20, RSV 3/12, Human rhnivoirus (RV) RV 16, or mock (MOI = 3). Additional details can be found in Supplemental Methods.

### NAEC DNAm acquisition and quality control

DNAm from NAECs was measured using the Asthma&Allergy (A&A) Custom BeadChip v1.0 and underwent quality control as previously described [8]. Additionally, probes overlapping common SNPs, with detection p-values *≥*0.01 in *≥*25% of samples, or on sex chromosomes were excluded. Outlier samples, identified by visual inspection of total methylation intensities and beta value box plots, were removed. Methylated and unmethylated intensities were quantile normalized and DNAm M-values were used.

### NAEC RNAseq

RNA sequencing (RNAseq) was available for 54 of the 88 NAECs samples collected at ages 2-3 years. Log_2_-transformed read counts per million were used in all analyses. Supplemental Methods contains additional details.

### Statistical analyses

We used the DNAm data to estimate the proportion of ciliated, suprabasal, and “other” cells via *in silico* cell type deconvolution (see Supplemental Methods for details). We identified RSV-associated CpGs at ages two and six years in each cell type by regressing DNAm onto the interaction between RSV infection by age 1 year (yes/no) and the cellular proportions defined above while adjusting for cellular proportions, daycare attendance during the first year of life (yes/no), older siblings (yes/no), sex, race, age of NAEC collection (two or six years), DNA concentration, DNAm plate, and 26 latent factors determined using CorrConf [9]. Individual-specific random effects were included. The regression coefficients for the six interaction terms (3 cell types *×* 2 ages) are the effects of infection on DNAm levels in each cell type, age pair [10]. Q-values [11] were used to control the false discovery rate at 5%.

We used linear mixed models to estimate the effect of infection on ALI cultured NAEC DNAm by regressing DNAm onto virus type (RSV 2014-1 01/2-20, RSV 3/12, RV 16, or mock) while adjusting for DNA concentration and four latent factors determined by CorrConf [9].

Because analyses involving gene expression data were performed in only 54 samples, we used an FDR threshold of 25% to identify genes whose expression was correlated with the DNAm at RSV-associated CpGs (Table 2) and genes predictive of wheeze risk (Fig. 2). Supplemental Methods contains additional analysis details.

**Figure 1:**
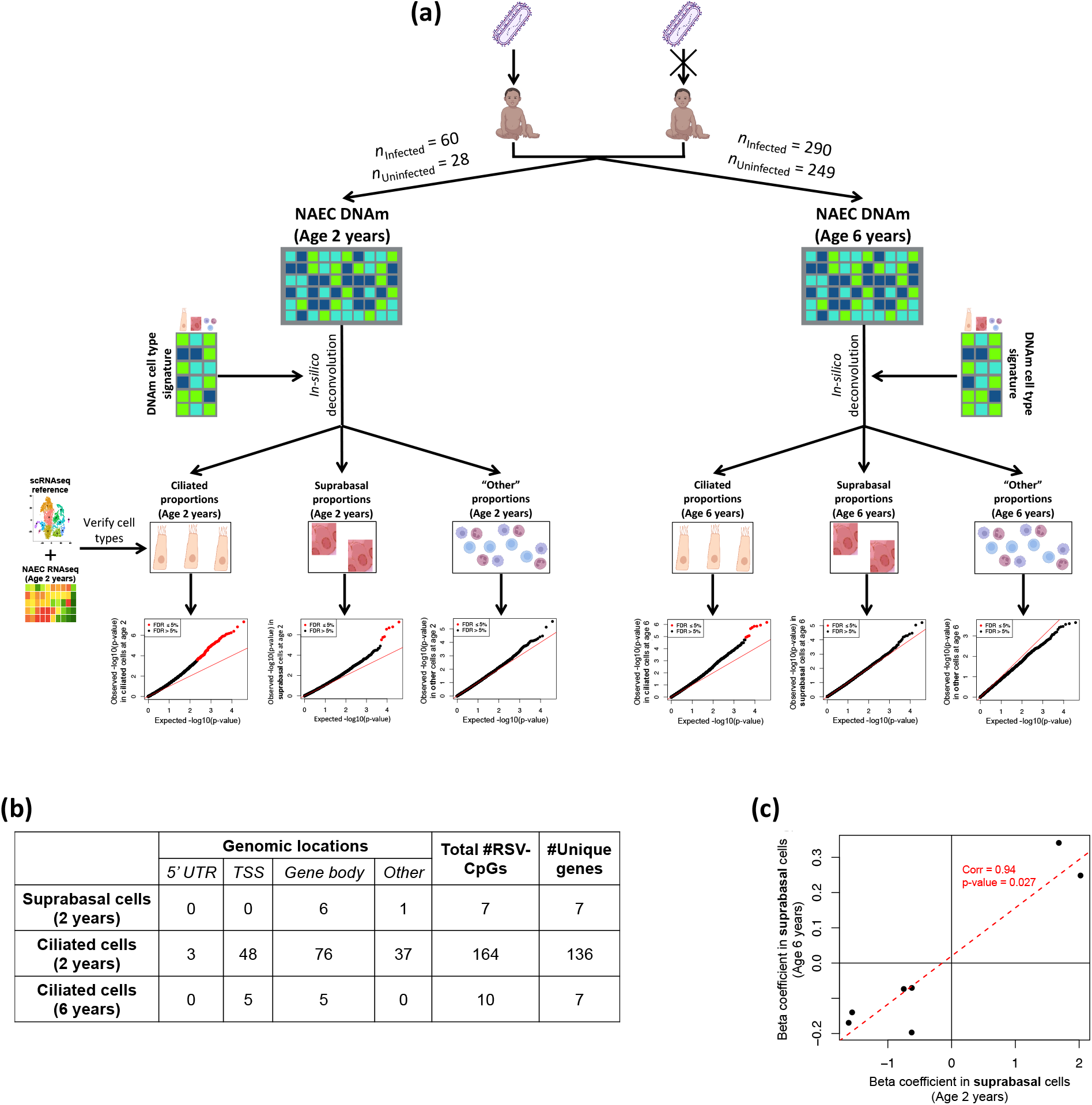
Determination of airway epithelial cell type-specific CpGs associated with infant (first year of life) RSV infection (RSV-CpGs). **(a)** Experimental design and analysis pipeline. NEAC DNAm data were deconvolved to estimate cell type proportions. Publicly available scRNAseq and bulk RNAseq collected from the same samples used to measure DNAm levels were used to confirm proportion estimates. Red points in the six Q-Q plots at the bottom of the figure panel ‘a’ indicate RSV-CpGs identified at age 2 years (left) and 6 years (right) in ciliated, suprabasal, and other cells (FDR ≤5%). **(b)** Genomic locations of RSV-CpGs. The sixth column (Total #RSV-CpGs) is the sum of the previous four columns; “TSS” is the *±* 2kb window around the transcriptional start site. **(c)** Beta regression coefficients for RSV-CpGs in suprabasal cells identified at age 2 years (x-axis) and their corresponding coefficient estimate at age 6 years (y-axis). The red line is the line of best fit. Abbreviations: 5’UTR, 5’ untranslated region; TSS, transcription start site

**Figure 2:**
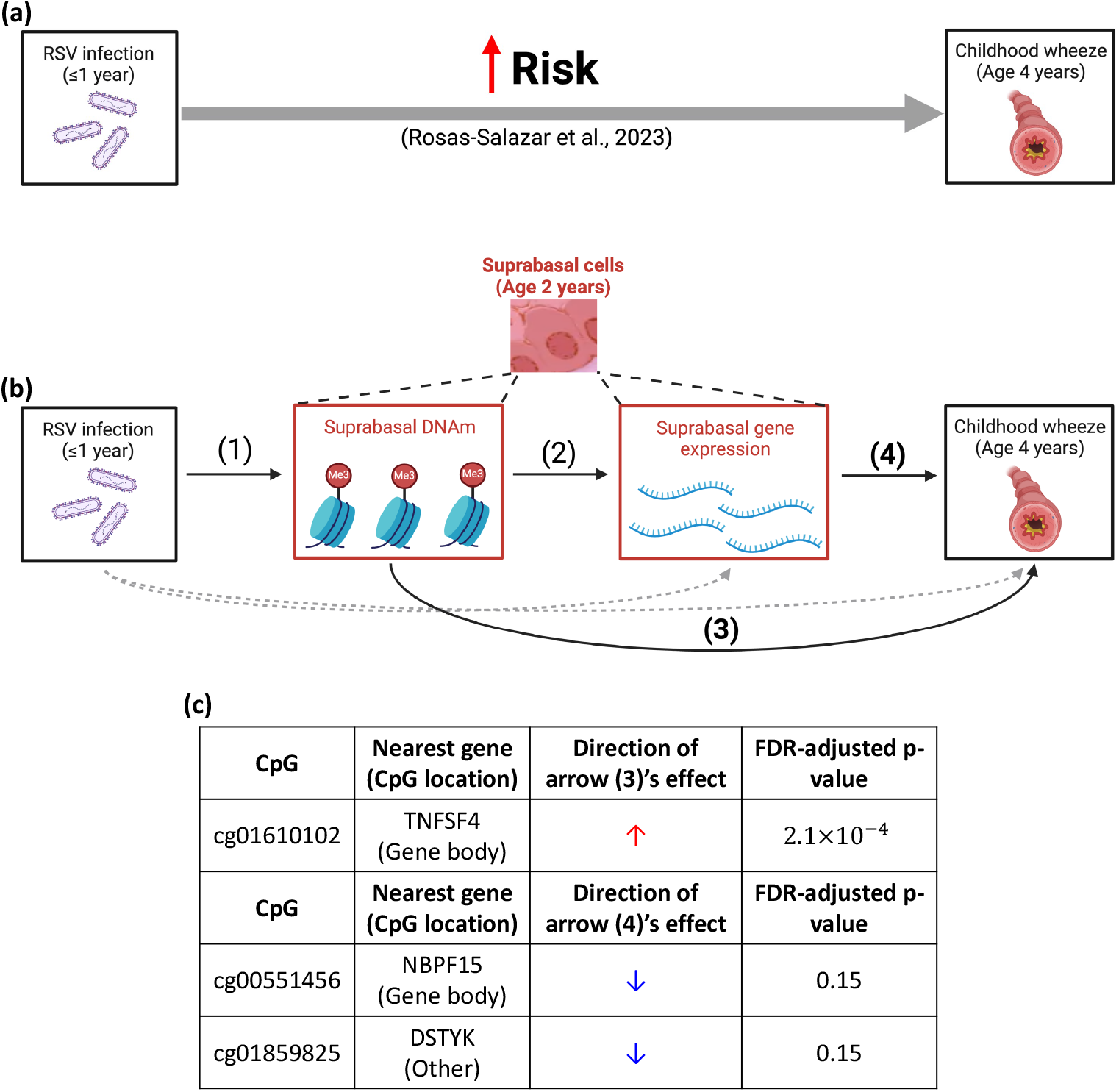
Assessment of the effect of first year RSV infection through DNAm on subsequent risk of wheeze **(a)**: The association of RSV infection and risk of developing childhood wheeze and asthma, based on results from Rosas-Salazar et al. [1]. **(b)**: Mechanism tested to explain the association of early-life RSV infection on wheeze and asthma. The direction of effects for arrows (1) and (2) are given in Table 2. Grey dashed arrows were not of interest, although we adjusted for them. **(c)**: P-values were adjusted for four tests (the number of CpGs and genes in Table 2).

## Results

### Population characteristics

Study participants were from the INSPIRE population-based birth cohort [12], where participant selection is described in the Methods. Characteristics for subjects participating in this study are given in Table 1. RSV infection status (not infected or infected) was ascertained in the first year of life using a combination of passive and active surveillance with viral identification through molecular and serological testing [1, Figure 2].

**Table 1:** Clinical and demographic information for study participants. The two age cohorts refer to the age at DNAm sampling.

|  | Age 2-3 years<br>(n = 88) | Age 6-7 years<br>(n = 539) |
| --- | --- | --- |
| Birth weight (g), median (IQR) | 3461.5 (603.2) | 3433.2 (567.4) |
| Gestational age (wk), median (IQR) | 39.0 (2.0) | 39.0 (1.5) |
| Sex |  |  |
| Male | 59.1% | 53.6% |
| Female | 40.9% | 46.4% |
| Race |  |  |
| White | 59.1% | 63.6% |
| Black | 23.9% | 18.9% |
| Hispanic | 4.5% | 9.1% |
| Other | 12.5% | 8.3% |
| Child breast-fed during the first year |  |  |
| Some breast-feeding | 79.5% | 84.8% |
| No breast-feeding | 20.5% | 15.2% |
| First-year daycare attendance |  |  |
| Yes | 36.3% | 37.1% |
| No | 63.7% | 62.9% |
| Does the child have older siblings? |  |  |
| Yes | 51.1% | 53.4% |
| No | 48.9% | 46.6% |
| Fourth-year wheeze |  |  |
| <2 wheezing episodes | 78.3% | 86.1% |
| ≥2 wheezing episodes | 21.7% | 13.9% |
| RSV infection status |  |  |
| Not infected by age 1 year | 31.9% | 46.2% |
| Infected by age 1 year | 68.1% | 53.8% |

### Inferring CpGs associated with infant RSV infection in each airway epithelial cell type

Since NAECs are a heterogeneous collection of cell types, we first used the DNAm data to estimate the proportion of ciliated, suprabasal, and “other” cell types in each NEAC sample, where we additionally leveraged our paired bulk RNA sequencing (RNAseq) data and publicly available single cell RNAseq (scRNAseq) data to verify proportion estimates (Supplemental Methods). The “other” cell type category captures non-ciliated and non-suprabasal cells and consists primarily of secretory cells (Supplemental Methods). Consistent with RSV-dependent changes in ciliated and secretory proportions observed *in vitro* [13], DNAm-derived ciliated and “other” proportions were lower and higher in children infected with RSV prior to age one year, respectively (p-values=0.025 and 0.041; Fig. S3). We used these proportions and the statistical method CellDMC [10] to estimate the effect of RSV infection on the DNAm levels in ciliated, suprabasal, and “other” cells (Methods). Briefly, CellDMC leverages the fact that bulk DNAm levels are a weighted average cell type-specific DNAm levels, where the weights are cell type proportions. We confirmed this was true in our NAEC DNAm data (Supplemental Methods). Consequently, the effect of RSV on each cell type’s DNAm can be estimated by regressing NAEC DNAm levels onto the interaction between cell type proportions and RSV infection status (yes/no). This method only relies on estimates for cell type proportions and does not require estimating cell type-specific DNAm levels. We define RSV-CpGs in each cell type to be those whose RSV regression coefficient was significant at a 5% false discover rate (FDR) in that cell type.

Of the 37,258 CpGs on the Asthma&Allergy array [8] that passed quality control, 164 and seven were RSV-CpGs at age two in ciliated and suprabasal cells, respectively (Fig. 1a-b). A dampened signal was observed at six years of age, where we identified only 10 RSV-CpGs in ciliated cells (Fig. 1a-b). Although no CpGs were genome-wide significant in suprabasal cells at age six, the beta coefficients at two and six years for the seven suprabasal RSV-CpGs identified at age two were highly correlated (corr=0.94, p-value=0.027, Fig. 1c; see Supplement for p-value calculation details), suggesting RSV-associated DNAm in suprabasal cells is temporally conserved. Such temporal consistency was not observed in ciliated cells. Table S1 provides the regression coefficients and p-values for RSV-CpGs in each NEAC type and age.

### Replication of DNAm changes following *in vitro* RSV infection in nasal airway epithelial cell cultures in air-liquid interface

Given that DNAm at the seven RSV-CpGs in suprabasal cells from children with RSV infection in infancy appears to be temporally conserved, we next sought to replicate this observation in an independent replication dataset derived from healthy infant NAECs grown in an air-liquid interface (ALI) medium and infected with two strains of RSV (see Methods). We did not observe statistically significant associations between *in vitro* RSV infection and the DNAm levels at individual RSV-CpGs after adjusting for 14 tests (2 strains *×* 7 RSV-CpGs), possibly because these replication data were derived from only eight donors. However, the directions of 11 out of the 14 replication beta coefficients matched those in our discovery dataset (p-value=0.050; see Supplement for calculation details; Table S2), providing further support for the associations between RSV infection and DNAm levels in suprabasal cells.

### Cell type-specific functional interpretation of RSV-CpGs

We next sought to determine the downstream functional relevance of RSV-CpGs by examining their impact on gene expression using paired NAEC DNAm and RNAseq data available for 54 study participants at 2 years of age. Given the cell type-specificity of RSV-CpGs and previous work showing that the correlation between DNAm and expression varies across cell types [14], we hypothesized that correlations between DNAm levels of RSV-CpGs and gene expression levels may also be cell type-specific. To address this, we utilized cell proportion estimates and the method developed in Cai et al. [15], which utilizes the same statistical model relating bulk and cell type-specific DNAm used to infer RSV-CpGs, to estimate the direction of the cell type-specific correlation between DNAm and gene expression (Supplemental Methods). We only estimate the direction of the correlation due to method constraints described in Supplemental Methods. Given the above replication results, we focused on the correlation in suprabasal cells.

Table 2 shows the relationships between DNAm at four of seven (57%) suprabasal RSV-CpGs that were correlated with expression of their nearest gene. This represents a 4.6-fold enrichment compared to randomly selected sets of seven CpGs from the Asthma&Allergy array (p-value=0.042, see Supplement for calculation details), suggesting that suprabasal RSV-CpGs are enriched for functionally relevant CpGs. The indirect effect direction in Table 2 is the implied direction of the DNAm-mediated effect of RSV infection on expression in suprabasal cells. DNAm at five of the 164 (3%) ciliated RSV-CpGs was also significantly correlated with expression (Table S4).

**Table 2:**
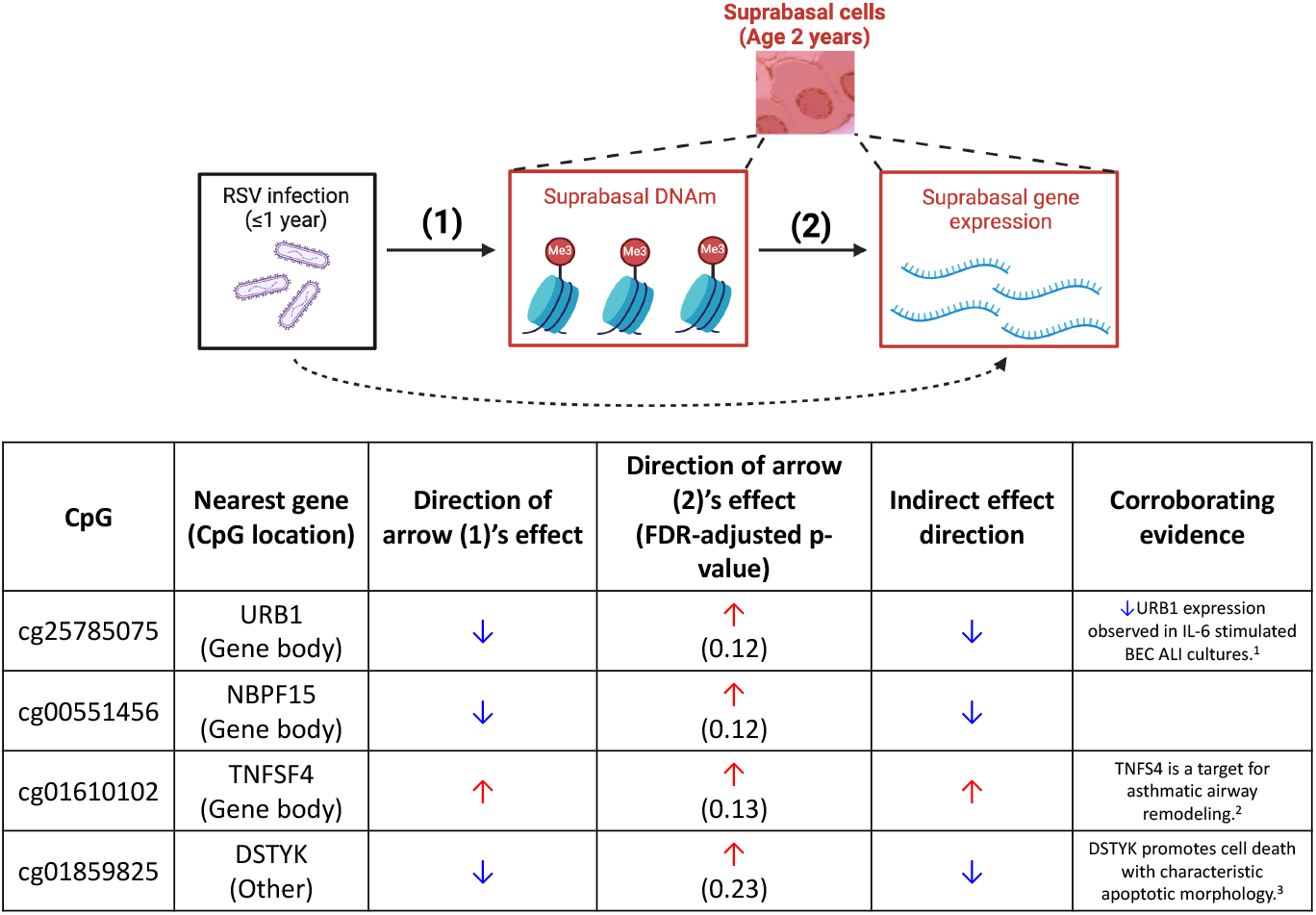
Results and functional interpretation for suprabasal RSV-CpGs whose DNAm was significantly correlated with its nearest gene’s expression in suprabasal cells, where the direction of arrow (2) is the direction of this correlation. The direction of arrow (1) was determined in “Inferring CpGs associated with infant RSV infection in each airway epithelial cell type”. The indirect effect is the product of the directions of arrows (1) and (2). We accounted for the direct effect (dashed line) in this analysis. ^1^: Jevnikar et al. [16]; ^2^: Seshasayee et al. [17]; ^3^: Zha et al. [18].

### Association of RSV-CpGs with childhood wheeze

Recently, Rosas-Salazar et al. [1] showed that RSV infection in the first year of life is associated with increased risk of wheezing and asthma later in childhood (Fig. 2a). We therefore sought to determine whether the RSV-CpGs we identified could provide a molecular mechanism by which RSV infection in the first year of life increases wheeze risk. Due to the temporal stability and robustness of suprabasal RSV-CpGs, we restricted our attention to the DNAm in suprabasal cells. We considered the mechanism in Fig. 2b which hypothesizes that the suprabasal DNAm at functionally relevant suprabasal RSV-CpGs (see Table 2), as well as the suprabasal expression of their corresponding genes, mediates the impact of infant RSV infection on subsequent risk of wheeze. The two indirect effects of interest in Fig. 2b are given by the two paths that are shown by solid arrows from RSV infection to childhood wheeze, namely, the paths given by arrows *{*(1), (2), (4)*}* and *{*(1), (3)*}*. The direction of an indirect effect is the product of the directions of effect for each solid arrow in the path. For example, if infection reduced a suprabasal DNAm at a CpG (arrow (1) is negative) and DNAm at that CpG in suprabasal cells was also inversely associated with wheeze risk (arrow (3) is negative), then the direction of the indirect effect would be such that infection increases wheeze risk. For the arrows describing the relationships, we focus on the direction of association and not the magnitudes due to method constraints described in Supplemental Methods. Due to sample size constraints, we did not have the power to explicitly test for mediation. Instead, since the directions of effects for arrows (1) and (2) were already estimated in Table 2, we estimated the direction of arrows (3) and (4) and used the resulting two indirect effects to interpret our results.

The estimated directions of arrows (3) and (4) for significant associations are given in Fig. 2c. While it was not statistically significant, the direction of arrow (4) for *TNFSF4* was also positive. Quite remarkably, the resulting estimated directions of the two indirect effects were positive for all CpGs in Figs. 2c, i.e., infection increases risk of wheeze. This is exactly what we would expect given the results of the Rosas-Salazar et al. study (Fig. 2a), and further suggests the RSV-CpGs we identify in suprabasal cells may mediate the impact of infant RSV infection on risk of subsequent childhood wheeze.

## Discussion

This study provides evidence to suggest that RSV infection in the first year of life has a lasting, cell type-specific impact on the DNAm in the airways of infants. We used *in vitro* RSV infection of nasal airway epithelial cells from healthy children differentiated in ALI cultured cells to replicate *ex vivo* observations of RSV-associated CpGs in suprabasal cells, and showed that these CpGs were enriched for CpGs correlated with suprabasal gene expression. We illustrated the clinical relevance of these CpGs and their associated genes by showing that their DNAm levels and expression of their nearest gene in suprabasal cells at age two years are predictive of recurrent wheeze risk at age four years.

early-life RSV infection is closely linked with the development of childhood wheeze and asthma, although there is considerable debate as to whether infection *causes* wheeze and asthma or if the association is confounded by host genetics or other environmental factors [19, 3]. Our study therefore sought to infer the impact of early-life RSV infection on airway DNAm later in childhood so as to identify potential molecular mechanisms through which infection leads to childhood wheeze and asthma. Importantly, we didn’t assess a clinical phenotype of severe RSV infection which likely identifies children with a shared genetic predisposition to both severe RSV and asthma [3]. Instead, we defined RSV infection in infancy as an *exposure*, determined using PCR or serological testing in healthy term infants. The prevalence of RSV infection and its independence of host genotype implies exposure is interpretable as a quasi-random event similar to “treatment assignment” in a randomized controlled trial [1, 20], which suggests that our analyses come closer to estimating the *causal* impact of infection on DNAm. Conversely, existing works exploring the association between RSV and DNAm define infection to be a severe manifestation of RSV infection such as RSV bronchiolitis [4, 5, 6], which is a clinical phenotype of RSV infection rather than an exposure. Associations in these studies may therefore reflect the myriad of confounding factors that impact severe RSV infection risk and DNAm, such as environmental exposures or host genetics [21, 22, 3].

We measured DNAm in NAECs instead of peripheral blood, the tissue of interest in existing studies of DNAm and RSV infection [4, 5, 6], because DNAm in NAECs is more strongly associated with asthma and allergy compared to studies in blood cells [23, 24]. We additionally used a novel Asthma&Allergy DNAm array enriched for asthma- and allergy-related CpGs to address the fact that CpGs on other commercial arrays were chosen without consideration of non-cancer diseases [8]. As NAECs are a heterogeneous mixture of cell types, we used cutting-edge statistical methods to deconvolve DNAm and gene expression data to infer cell type-specific effects. This represents a marked departure from existing epigenome-wide analyses that implicitly assume the effects of interest are the same across cell types.

We identified seven and 164 RSV-CpGs in suprabasal and ciliated cells, respectively, at age two years. We observed more RSV-associated DNAm in ciliated cells possibly because NAECs contain more ciliated than suprabasal cells (Fig. S3), meaning we have more power to identify RSV-CpGs in ciliated cells. A second possibility is that RSV primarily infects ciliated cells [25]. While ciliated cells are a terminally differentiated cell type [26], they may survive for long periods in early-life, with one study estimating their half-life in the trachea and lung of mice to be six and 17 months, respectively. We note, however, there are no comparable studies in humans of which we are aware [26]. Thus it is possible that some of the ciliated cells we captured at age two years may have been infected with RSV before age one year, where these cells may harbor more RSV-induced DNAm. While the impact of infection on DNAm was dampened at age six years, likely due to the large difference between time of infection and DNAm measurement, there was a remarkable and statistically significant concordance in RSV-associated DNAm in suprabasal cells at ages two and six years, suggesting infection has a lasting impact on the DNAm in suprabasal cells. This RSV signature in suprabasal DNAm was further validated using *in vitro* RSV infection in ALI cultures.

The enduring impact of infection suggests RSV infection in early-life has a long-lasting impact on the epigenetic programs of basal-related cells in the airway. This “epigenetic memory” of airway stem cells is congruent with other recent experimental results [27]. For example, Ordovas-Montanes et al. [28] showed that the levels of the Wnt pathway activator *CTNNB1* in ALI cultures of healthy NAEC-derived basal cells exposed to Type 2 cytokines resemble *CTNNB1* levels from unexposed polyp-derived basal cells. It is also consistent with work from our group showing that ALI cultured NAECs of childhood wheezers who were infected with RSV in early-life contained basal cells with altered differentiation trajectories [29]. The functional relevance of RSV-induced changes to non-ciliated cells in general and basal cells in particular has also been observed in rhinovirus infection [30, 31].

The correlation between the DNAm at suprabasal RSV-CpGs and the expression of their nearest genes allowed us to assess the functionality of suprabasal RSV-CpGs by determining how they mediated the impact of RSV infection on expression (Table 2). The impact of infection on the DNAm of *URB1* has the effect of decreasing the expression of this gene in suprabasal cells, which is consistent with studies showing that the expression of *URB1* in bronchial epithelial cells is reduced upon stimulation with IL-6, a cytokine released into the airway upon RSV infection [16, 32]. Reductions in URB1 protein levels impair ribosome biogenesis [33], which in turn has been shown to reduce type I interferon production [34], where lower levels of these cytokines are often observed in children with recurrent wheeze [35]. We also observed that the suprabasal expression of *TNFSF4* (i.e., *OX40L*), which encodes the ligand for the T-cell activation marker TNFRSF4, is up-regulated, which is consistent with its role as a facilitator of Type 2 inflammation in asthmatics [17]. The reduction in the expression of *DSTYK* is consistent with its role as a promoter of apoptosis [36], as prolonged or disrupted apoptosis is characteristic of asthma phenotypes [37, 38, 39].

We lastly showed that suprabasal RSV-CpGs and their mapped genes may mediate the effect of early-life RSV infection on childhood wheeze (Fig. 2). This not only indicates that these CpGs are clinically relevant, but also provides a possible molecular mechanism through which early-life RSV infection *causes* subsequent childhood wheeze and respiratory morbidity [1]. These results provide additional evidence that DNAm mediates the impact of early-life exposures [40] and further supports the hypothesis that epigenetic changes to airway stem cells are relevant to asthma-related disorders [27].

Despite our favorable study design, robust results, and the biological plausibility of our findings, we must acknowledge some limitations. First, we did not have NAEC samples for DNAm at a timepoint prior to the first RSV infection. Seond, we used computational methods to deconvolve bulk DNAm and gene expression data and infer cell type-specific associations. While these methods are widely used and give reliable estimates [10, 41], our results should be confirmed using DNAm and expression from sorted or single cells. Third, since NAEC brushings contain few basal cells [42], we could only infer associations in suprabasal cells. While it is likely that the RSV-dependent epigenetic changes we observed in suprabasal cells also exist in basal cells, additional experiments are needed to confirm this. Fourth, we only used eight donors in our ALI replication experiments, which limited our power to replicate individual RSV-CpGs. However, the directions of the associations between RSV and DNAm levels in our replication experiments matched what we observed in suprabasal cells, providing additional evidence that the RSV-CpGs identified in suprabasal cells were legitimate.

In summary, using a human birth cohort and *in vitro* RSV infection we identified seven novel CpGs whose DNAm levels in suprabasal airway epithelial cells were associated with infant RSV infection. Our results reveal that these epigenetic changes in suprabasal cells of children who have had RSV infection in the first year of life are conserved over time, associated with subsequent wheeze, have functional relevance, and can be replicated in *in vitro* infection in ALI cultures. These findings help us posit a molecular mechanism through which early-life RSV infection may lead to childhood respiratory morbidity and asthma.

## Supporting information

Supplementary analysis details

Regression estimates for all RSV-CpGs in each cell type and age cohort

ALI replication results

## Acknowledgements

We are deeply grateful to all of the families who participated in this study, to the INSPIRE research study staff and to the middle Tennessee pediatric practices with whom we collaborated to enroll a representative population of our region.

## Declaration of funding sources

This work was supported by the CREW consortium, Office of the Director, National Institutes of Health under Award Number UG3/UH3OD023282 and the National Institute of Allergy and Infectious Diseases under Award Number U19 AI62310. Additional funding that supported this research came from National Institute of Allergy and Infectious Diseases (under award number K24AI77930); The National Institute on Aging (R01AG080590-01); the Vanderbilt Institute for Clinical and Translational Research (the National Center for Advancing Translational Sciences under award number UL1TR000445) and UM1 AI160040, UG3 OD035509. The content is solely the responsibility of the authors and does not necessarily represent the official views of the funding agencies. The study sponsors had no role in the study design; in the collection, analysis, and interpretation of data; in the writing of the report; and in the decision to submit the paper for publication. The authors were not paid to write this article by a pharmaceutical company or other agency.

## Conflict of interest

Dr. McKennan reported grants from NIH during the conduct of the study and personal fees from SignatureDx outside the submitted work. Dr. Bacharier reported grants from NIH/National Institute of Allergy and Infectious Diseases (NIAID) during the conduct of the study and personal fees from GlaxoSmithKline, Genentech/Novartis, Merck, DBV Technologies, Teva, Boehringer Ingelheim, AstraZeneca, Sanofi/Regenerol. Dr. Gern reported grants from NIH during the conduct of the study, personal fees from AstraZeneca and Meissa Vaccines Inc., and stock options for Meissa Vaccines Inc. outside the submitted work. Dr. Hartert reported grants from NIH and the World Health Organization during the conduct of the study and personal fees from the American Thoracic Society, the NIH, Parker B. Francis Foundation, Pfizer and Sanofi-Pasteur outside the submitted work.

## Authors’ Contributions

CM, TH, TG and DN contributed to the conception of the study and/or the study design. TH, CM, SB, JG and DN obtained the research funding supporting this study. TH designed and conducted the clinical study, biospecimen acquisition, supervision and project administration. TG contributed to the study design, derivation of clinical variables and power analyses. DN led the nasal airway epithelial culture, cryopreservation and *in vitro* experiments. CO designed the DNA mehtylation array. Data analyses were designed and performed by CM. The manuscript and figures were drafted by CM and TH. All authors read and approved the final manuscript.

## References

[1] C. Rosas-Salazar, T. Chirkova, T. Gebretsadik, et al. “Respiratory syncytial virus infection during infancy and asthma during childhood in the USA (INSPIRE): a population-based, prospective birth cohort study”. In: The Lancet 401.10389 (May 2023), pp. 1669–1680. doi: 10.1016/s0140-6736(23)00811-5. URL: 10.1016/s0140-6736(23)00811-5.

[2] J. G. Wildenbeest, M.-N. Billard, R. P. Zuurbier, et al. “The burden of respiratory syncytial virus in healthy term-born infants in Europe: a prospective birth cohort study”. In: The Lancet Respiratory Medicine 11.4 (Apr. 2023), pp. 341–353. doi: 10.1016/s2213-2600(22)00414-3. URL: 10.1016/s2213-2600(22)00414-3.

[3] S. M. Brunwasser, B. M. Snyder, A. J. Driscoll, et al. “Assessing the strength of evidence for a causal effect of respiratory syncytial virus lower respiratory tract infections on subsequent wheezing illness: a systematic review and meta-analysis”. In: The Lancet Respiratory Medicine 8.8 (Aug. 2020), pp. 795–806. doi: 10.1016/s2213-2600(20)30109-0. URL: 10.1016/s2213-2600(20)30109-0.

[4] S. Pischedda, I. Rivero-Calle, A. Gómez-Carballa, et al. “Role and Diagnostic Performance of Host Epigenome in Respiratory Morbidity after RSV Infection: The EPIRESVi Study”. In: Frontiers in Immunology 13 (May 2022). ISSN: 1664-3224. doi: 10.3389/fimmu.2022.875691. URL: 10.3389/fimmu.2022.875691.

[5] M. Elgizouli, C. Logan, R. Grychtol, et al. “Reduced PRF1 enhancer methylation in children with a history of severe RSV bronchiolitis in infancy: an association study”. In: BMC Pediatrics 17.1 (Mar. 2017). ISSN: 1471-2431. doi: 10.1186/s12887-017-0817-9. URL: 10.1186/s12887-017-0817-9.

[6] Z. Zhu, Y. Li, R. J. Freishtat, et al. “Epigenome-wide association analysis of infant bronchiolitis severity: a multicenter prospective cohort study”. In: Nature Communications 14.1 (Sept. 2023). ISSN: 2041-1723. doi: 10.1038/s41467-023-41300-y. URL: 10.1038/s41467-023-41300-y.

[7] M. M. Soliai, A. Kato, K. A. Naughton, et al. “Epigenetic responses to rhinovirus exposure in airway epithelial cells are correlated with key transcriptional pathways in chronic rhinosinusitis”. In: Allergy 78.10 (Aug. 2023), pp. 2698–2711. ISSN: 1398-9995. doi: 10.1111/all.15837.

[8] A. Morin, E. E. Thompson, B. A. Helling, et al. “A functional genomics pipeline to identify high-value asthma and allergy CpGs in the human methylome”. In: Journal of Allergy and Clinical Immunology 151.6 (June 2023), pp. 1609–1621. doi: 10.1016/j.jaci.2022.12.828.

[9] C. McKennan and D. Nicolae. “Estimating and Accounting for Unobserved Covariates in High-Dimensional Correlated Data”. In: Journal of the American Statistical Association 117.537 (June 2020), pp. 225–236. doi: 10.1080/01621459.2020.1769635. URL: 10.1080/01621459.2020.1769635.

[10] S. C. Zheng, C. E. Breeze, S. Beck, et al. “Identification of differentially methylated cell types in epigenome-wide association studies”. In: Nature Methods 15.12 (2018), pp. 1059–1066.

[11] J. D. Storey. “A Direct Approach to False Discovery Rates”. In: Journal of the Royal Statistical Society Series B: Statistical Methodology 64.3 (Aug. 2002), pp. 479–498. doi: 10.1111/1467-9868.00346.

[12] E. K. Larkin, T. Gebretsadik, M. L. Moore, et al. “Objectives, design and enrollment results from the Infant Susceptibility to Pulmonary Infections and Asthma Following RSV Exposure Study (INSPIRE)”. In: BMC Pulmonary Medicine 15.1 (Apr. 2015). doi: 10.1186/s12890-015-0040-0. URL: 10.1186/s12890-015-0040-0.

[13] B. D. Persson, A. B. Jaffe, R. Fearns, et al. “Respiratory Syncytial Virus Can Infect Basal Cells and Alter Human Airway Epithelial Differentiation”. In: PLoS ONE 9.7 (July 2014). Ed. by K. Harrod, e102368. doi: 10.1371/journal.pone.0102368. URL: 10.1371/journal.pone.0102368.

[14] K. Wang, R. Dai, Y. Xia, et al. “Spatiotemporal specificity of correlated DNA methylation and gene expression pairs across different human tissues and stages of brain development”. In: Epigenetics 17.10 (Oct. 2021), pp. 1110–1127. doi: 10.1080/15592294.2021.1993607. URL: 10.1080/15592294.2021.1993607.

[15] B. Cai, J. Zhang, H. Li, et al. Statistical Inference of Cell-type Proportions Estimated from Bulk Expression Data. 2022. arXiv: 2209.04038 [stat.ME].

[16] Z. Jevnikar, J. Östling, E. Ax, et al. “Epithelial IL-6 trans-signaling defines a new asthma phenotype with increased airway inflammation”. In: Journal of Allergy and Clinical Immunology 143.2 (Feb. 2019), pp. 577–590. doi: 10.1016/j.jaci.2018.05.026. URL: 10.1016/j.jaci.2018.05.026.

[17] D. Seshasayee, W. P. Lee, M. Zhou, et al. “In vivo blockade of OX40 ligand inhibits thymic stromal lymphopoietin driven atopic inflammation”. In: Journal of Clinical Investigation 117.12 (Dec. 2007), pp. 3868–3878. ISSN: 0021-9738. doi: 10.1172/jci33559. URL: 10.1172/JCI33559.

[18] J. Zha, Q. Zhou, L.-G. Xu, et al. “RIP5 is a RIP-homologous inducer of cell death”. In: Biochemical and Biophysical Research Communications 319.2 (June 2004), pp. 298–303. doi: 10.1016/j.bbrc.2004.04.194. URL: 10.1016/j.bbrc.2004. 04.194.

[19] T. Jartti and J. E. Gern. “Role of viral infections in the development and exacerbation of asthma in children”. In: Journal of Allergy and Clinical Immunology 140.4 (Oct. 2017), pp. 895–906. ISSN: 0091-6749. doi: 10.1016/j.jaci.2017.08.003. URL: 10.1016/j.jaci.2017.08.003.

[20] D. Lawless, C. G. McKennan, S. R. Das, et al. “Viral Genetic Determinants of Prolonged Respiratory Syncytial Virus Infection Among Infants in a Healthy Term Birth Cohort”. In: The Journal of Infectious Diseases 227.10 (Nov. 2022), pp. 1194–1202. doi: 10.1093/infdis/jiac442. URL: 10.1093/infdis/jiac442.

[21] M. H. Lee, D. Mailepessov, K. Yahya, et al. “Air quality, meteorological variability and pediatric respiratory syncytial virus infections in Singapore”. In: Scientific Reports 13.1 (Jan. 2023). ISSN: 2045-2322. doi: 10.1038/s41598-022-26184-0. URL: 10.1038/s41598-022-26184-0.

[22] C. Lau, J. C. Behlen, A. Myers, et al. “In Utero Ultrafine Particulate Exposure Yields Sex-and Dose-Specific Responses to Neonatal Respiratory Syncytial Virus Infection”. In: Environmental Science & Technology 56.16 (2022), pp. 11527–11535. doi: 10.1021/acs.est.2c02786.

[23] M. van Breugel, C. Qi, Z. Xu, et al. “Nasal DNA methylation at three CpG sites predicts childhood allergic disease”. In: Nature Communications 13.1 (Dec. 2022). ISSN: 2041-1723. doi: 10.1038/s41467-022-35088-6. URL: 10.1038/s41467-022-35088-6.

[24] C. Qi, Y. Jiang, I. V. Yang, et al. “Nasal DNA methylation profiling of asthma and rhinitis”. In: Journal of Allergy and Clinical Immunology 145.6 (June 2020), pp. 1655–1663. ISSN: 0091-6749. doi: 10.1016/j.jaci.2019.12.911. URL: 10.1016/j.jaci.2019.12.911.

[25] L. Zhang, M. E. Peeples, R. C. Boucher, et al. “Respiratory Syncytial Virus Infection of Human Airway Epithelial Cells Is Polarized, Specific to Ciliated Cells, and without Obvious Cytopathology”. In: Journal of Virology 76.11 (June 2002), pp. 5654–5666. doi: 10.1128/jvi.76.11.5654-5666.2002. URL: 10.1128/jvi.76.11.5654-5666.2002.

[26] E. L. Rawlins and B. L. M. Hogan. “Ciliated epithelial cell lifespan in the mouse trachea and lung”. In: American Journal of Physiology-Lung Cellular and Molecular Physiology 295.1 (2008). PMID: 18487354, pp. L231–L234. doi: 10.1152/ajplung.90209.2008. eprint: 10.1152/ajplung.90209.2008. URL: 10.1152/ajplung.90209.2008.

[27] E. Ruysseveldt, K. Martens, and B. Steelant. “Airway Basal Cells, Protectors of Epithelial Walls in Health and Respiratory Diseases”. In: Frontiers in Allergy 2 (Nov. 2021). ISSN: 2673-6101. doi: 10.3389/falgy.2021.787128. URL: 10.3389/falgy.2021.787128.

[28] J. Ordovas-Montanes, D. F. Dwyer, S. K. Nyquist, et al. “Allergic inflammatory memory in human respiratory epithelial progenitor cells”. In: Nature 560.7720 (Aug. 2018), pp. 649–654. ISSN: 1476-4687. doi: 10.1038/s41586-018-0449-8. URL: 10.1038/s41586-018-0449-8.

[29] S. Berdnikovs, D. C. Newcomb, K. E. McKernan, et al. “Single cell profiling to determine influence of wheeze and early-life viral infection on developmental programming of airway epithelium”. In: bioRxiv (2024). doi: 10.1101/2024.07.08.602506. eprint: https://www.biorxiv.org/content/early/2024/07/11/2024.07.08.602506.full.pdf. URL: https://www.biorxiv.org/content/early/2024/07/11/2024.07.08.602506.

[30] J. Nicodemus-Johnson, K. A. Naughton, J. Sudi, et al. “Genome-Wide Methylation Study Identifies an IL-13–induced Epigenetic Signature in Asthmatic Airways”. In: American Journal of Respiratory and Critical Care Medicine 193.4 (Feb. 2016), pp. 376–385. ISSN: 1535-4970. doi: 10.1164/rccm.201506-1243oc. URL: 10.1164/rccm.201506-1243OC.

[31] S. Djeddi, D. Fernandez-Salinas, G. X. Huang, et al. “Rhinovirus infection of airway epithelial cells uncovers the non-ciliated subset as a likely driver of genetic susceptibility to childhood-onset asthma”. In: medRxiv (Feb. 2024). doi: 10.1101/2024.02.02.24302068. eprint: https://www.medrxiv.org/content/early/2024/02/06/2024.02.02.24302068.full.pdf. URL: https://www.medrxiv.org/content/early/2024/02/06/2024.02.02.24302068.

[32] C. J. Pyle, F. I. Uwadiae, D. P. Swieboda, et al. “Early IL-6 signalling promotes IL-27 dependent maturation of regulatory T cells in the lungs and resolution of viral immunopathology”. In: PLOS Pathogens 13.9 (Sept. 2017). Ed. by C. B. Lopez, e1006640. ISSN: 1553-7374. doi: 10.1371/journal.ppat.1006640. URL: 10.1371/journal.ppat.1006640.

[33] L. Shan, G. Xu, R.-W. Yao, et al. “Nucleolar URB1 ensures 3 ETS rRNA removal to prevent exosome surveillance”. In: Nature 615.7952 (Mar. 2023), pp. 526–534. ISSN: 1476-4687. doi: 10.1038/s41586-023-05767-5. URL: 10.1038/s41586-023-05767-5.

[34] C. Bianco and I. Mohr. “Ribosome biogenesis restricts innate immune responses to virus infection and DNA”. In: eLife 8 (Dec. 2019). ISSN: 2050-084X. doi: 10.7554/elife.49551. URL: 10.7554/eLife.49551.

[35] H. Makrinioti, A. Bush, J. Gern, et al. “The Role of Interferons in Driving Susceptibility to Asthma Following Bronchiolitis: Controversies and Research Gaps”. In: Frontiers in Immunology 12 (Dec. 2021). ISSN: 1664-3224. doi: 10.3389/fimmu.2021.761660. URL: 10.3389/fimmu.2021.761660.

[36] J. Zha, Q. Zhou, L.-G. Xu, et al. “RIP5 is a RIP-homologous inducer of cell death”. In: Biochemical and Biophysical Research Communications 319.2 (June 2004), pp. 298–303. doi: 10.1016/j.bbrc.2004.04.194. URL: 10.1016/j.bbrc.2004. 04.194.

[37] T. P. Moran and B. P. Vickery. “Apoptotic Cell Clearance by Bronchial Epithelial Cells Critically Influences Airway Inflammation”. In: Pediatrics 132.Supplement1 (Oct. 2013), S37–S38. ISSN: 1098-4275. doi: 10.1542/peds.2013-2294jjj. URL: 10.1542/peds.2013-2294JJJ.

[38] H. Yamaguchi, T. Maruyama, Y. Urade, et al. “Immunosuppression via adenosine receptor activation by adenosine monophosphate released from apoptotic cells”. In: eLife 3 (Mar. 2014). Ed. by D. Wallach, e02172. ISSN: 2050-084X. doi: 10.7554/eLife.02172. URL: 10.7554/eLife.02172.

[39] L. Liu, L. Zhou, L.-L. Wang, et al. “Programmed Cell Death in Asthma: Apoptosis, Autophagy, Pyroptosis, Ferroptosis, and Necroptosis”. In: Journal of Inflammation Research Volume 16 (July 2023), pp. 2727–2754. ISSN: 1178-7031. doi: 10.2147/jir.s417801. URL: 10.2147/JIR.S417801.

[40] C. Mitchell, L. M. Schneper, and D. A. Notterman. “DNA methylation, early life environment, and health outcomes”. In: Pediatric Research 79.1–2 (Oct. 2015), pp. 212–219. ISSN: 1530-0447. doi: 10.1038/pr.2015.193. URL: 10.1038/pr.2015.193.

[41] M. Cai, J. Zhou, C. McKennan, et al. “scMD facilitates cell type deconvolution using single-cell DNA methylation references”. In: Communications Biology 7.1 (Jan. 2024). ISSN: 2399-3642. doi: 10.1038/s42003-023-05690-5. URL: 10.1038/s42003-023-05690-5.

[42] M. Deprez, L.-E. Zaragosi, M. Truchi, et al. “A Single-Cell Atlas of the Human Healthy Airways”. In: American Journal of Respiratory and Critical Care Medicine 202.12 (Dec. 2020), pp. 1636–1645. doi: 10.1164/rccm.201911-2199oc. URL: 10.1164/rccm.201911-2199oc.

