## Supplementary analysis details for "Impact of early-life respiratory syncytial virus infection on cell type-specific airway DNA methylation"

September 29, 2024

#### S1 Overview

This supplement contains additional supplemental figures, tables, and methods. Table S1 ("TableS1.xlsx") contains the regression estimates for all RSV-CpGs in each cell type and age cohort, and Table S2 ("TableS2.xlsx") contains the ALI replication results.

#### S2 Supplemental non-statistical methods

##### S2.1 NAEC RNAseq

After passing through the QIAshredder, RNA was extracted using the RNeasy Mini Kit (Qiagen, Hilden, Germany). RNA quality was assessed using a NanoDrop 2000 Spectrophotometer (Thermo Fisher Scientific, Waltham, USA). cDNA was generated using 125.4 ug of total RNA and the SuperScript IV First-Strand Synthesis kit (Thermo Fisher Scientific, Waltham, USA). qPCR was conducted for IFNL2, RNA M gene, TJP1, KRT5, and ALDOA using QuantStudio. IFNL2 primers (Hs00820125_g1) were purchased from Applied Biosystems and the RNA M gene (forward: GGC AAA TAT GGA AAC ATA CGT GAA; reverse: TCT TTT TCT AGG ACA TTG TAY TGA ACA) was generated using IDT. TJP1 (Hs01551861_m1), KRT5 (Hs00361185_m1), and ALDOA (Hs00605108_g1) primers were purchased from Applied Biosystems.

##### S2.2 Airway epithelial culture

NAECs collected at ages 2-3 years were placed in PneumaCult-Ex Plus Medium (StemCell, Vancouver, Canada). Cells were placed on collagen coated flasks and were submerged and expanded in PneumaCult-Ex Plus Medium, consisting of 38.76 ml of Ex Plus Basal Medium supplemented with 1.25 aliquots of PneumaCult-Ex Plus supplements. PneumaCult-Ex Plus supplements were a mix of 50 mL PneumaCult-Ex Plus 50X supplement, 2.5 mL

Hydrocortisone stock solution, and 5 mL Pen-Strep. Once cells reached 70-80% confluence on the flask, NAECs were disassociated using 2 ml Animal Component-Free (ACF) Cell Dissociation Solution and 2 ml ACF Inhibition Solution (StemCell, Vancouver, Canada) and transferred to a 24 transwell plate with transwells coated in 200 µl of collagen IV. Cells remained submerged in PneumaCult-Ex Plus Medium until confluency (approximately 1 week) with media changed every other day. Once confluent, media was removed from the top chamber and NAECs were allowed to differentiate for 3 weeks using PneumaCult ALI Medium (1ml) in the bottom chamber. Media in the bottom chamber was changed every other day. Complete PneumaCult ALI Medium consists of 35 mL of PneumaCult ALI medium and a 5 mL of ALI supplement aliquot.

##### S2.3 NAEC infection and harvest

Cells infected with RSV 2014-1 01/2-20, RSV 3/12, and mock were incubated for an hour at 37°C with gentle rocking, and cells infected with RV 16 were incubated while gently rocking for an hour at 33°C. After an hour, inoculum was removed from the apical surface, and cells were placed back into the appropriate temperature incubator. After 24 hours, 100uL of basolateral supernatants for each treatment were collected. After 48 hours, 200 µL of basolateral supernatants were collected for each treatment, and cells were harvested from transwells for RNA by adding 150 µL buffer RLT+ to the membrane for 15 minutes per the RNeasy Mini Kit (Qiagen, Hilden, Germany). Lysed cells were passed through a QIAshredder (Qiagen, Hilden, Germany).

#### S3 Supplemental statistical methods

##### S3.1 Estimates of cell type proportions

A key step in our analysis pipeline (Fig. 1a) is deconvolving bulk nasal airway epithelial cell (NAEC) DNAm to recover cell proportions. We describe our pipeline below, which involved first deconvolving DNAm into three cell types and then using bulk RNA sequencing collected in a subset of study participants, publicly available single cell RNA sequencing data, and well-defined cell type marker genes to properly label the aforementioned three cell types.

We deconvolved our NAEC DNAm data by constructing the first NAEC cell type signature matrix for CpGs on the Asthma and Allergy (A&A) array using NAEC DNAm and technician-labeled "ciliated", "squamous", and "other" cell counts based on microscopy as reported in Morin et al. [1]. Briefly, NAEC DNAm and available ciliated, squamous, and other cell counts from Morin et al. were first used to construct a cell type signature matrix for CpGs on the A&A array as the coefficients from the regression of DNAm M-values onto observed cellular proportions. Section S3.2 shows that this signature is appropriate for our DNAm data. We then estimated ciliated, squamous, and other cell proportions in our data by regressing bulk DNAm onto the aforementioned signature matrix via non-negative least squares with a sum-to-one constraint.

Contrary to the above technician-labeled cell types, the airway single-cell atlas developed by Deprez et al. [2] showed that nasal brushings are composed almost entirely of suprabasal, ciliated, secretory, and immune cells. Given that these samples were collected by trained research nurses at the inferior turbinate, we first hypothesized that technician-labeled "squamous" cells were predominantly suprabasal cells. To test this, we used single cell RNAseq (scRNAseq) data collected from the nasal brushings of healthy individuals in Ziegler et al. [3] to build a gene expression signature matrix for 11 NAEC types (Figure S1(a)). The suprabasal marker genes identified in Deprez et al. [2], which distinguish suprabasal from all other cell types found in nasal brushings, were used to identify cell type cluster 5 as the suprabasal cluster (Figure S1(b)). Notably, the two suprabasal marker genes in S1(b) are either not expressed or are expressed at trivial levels in squamous cells [4]. We then used MuSiC [5] to estimate cell type proportions for the aforementioned 11 cell types in the subset of INSPIRE participants that also had bulk RNAseq measurements. Consistent with our hypothesis, Figure S1(c) shows that DNAm-derived "squamous" cell proportions are highly correlated with suprabasal proportions (p-value $= 5.02 \times 10^{-6}$). Further, Figure S2 shows that DNAm-derived "squamous" cell proportions are unrelated to the proportions of the remaining 10 scRNAseq-derived cell types, where we used Lin's concordance correlation coefficient (CCC) [6] to quantify the concordance between proportions. The CCC takes values between -1 and 1 and measures the deviation from the red line $y = x$, where value closer to 1 mean points lie closer to the line $y = x$. It is preferable to Pearson correlation in this setting, since the correlation only measures the deviation from the line of best fit, which may not be $y = x$. Taken together, these results indicate that squamous proportion estimates are actually estimates for suprabasal proportions.

We next sought to confirm that DNAm-derived ciliated proportions are in fact ciliated proportions and identify the composition of "other" cell types. To do so, we first note that MuSiC estimated that in addition to cluster 5 in Figure S1(a) (the suprabasal clusters), clusters 1, 6, 8, 9, 10, and 11 were present with non-zero proportions in at least one sample. Since cluster 10's non-zero proportions were trivially small (Figure S2), we did not consider it in subsequent analyses. To infer the identities of these clusters, we first defined their marker genes as genes whose FDR-adjusted Wilcoxin rank p-values comparing their expression in that cluster to their expression in all other clusters were $\leq 0.05$. We then overlapped these markers with the ciliated, immune, and secretory cell markers identified in Vieira Braga et al. [7] and Ziegler et al. [3] to label them as ciliated, immune, or secretory cell clusters.

We first found that cluster 1, as well as clusters 0 and 2, were unequivocally a cluster of ciliated cells, since 28 out of the 31 ciliated marker genes were among cluster 1's markers and only three ciliated markers were among clusters 3-11's markers. Cluster 6 was a secretory cluster, since it was the only cluster whose markers included MUC5AC. Cluster 8 was likely another secretory cluster and clusters 9 and 11 were likely immune related clusters based on their markers' overlap with the secretory and immune markers identified in Ziegler et al. [3].

Next, we found that the correlation between DNAm-derived ciliated proportions and RNAseq-derived cluster 1's proportions (the ciliated cluster) was 0.31 (p-value = 0.012), suggesting DNAm-derived ciliated proportions are in fact ciliated proportions. We also found that the correlation between DNAm-derived "other" proportions and the remaining RNAseq-derived proportions (i.e. clusters 6, 8, 9, and 11) was 0.23 (p-value = 0.050), which suggests that "other" cells are likely a mixture of secretory and immune cells. Notably, secretory cells accounted for more than 65% of these remaining proportions, indicating "other" cells are predominantly secretory cells.

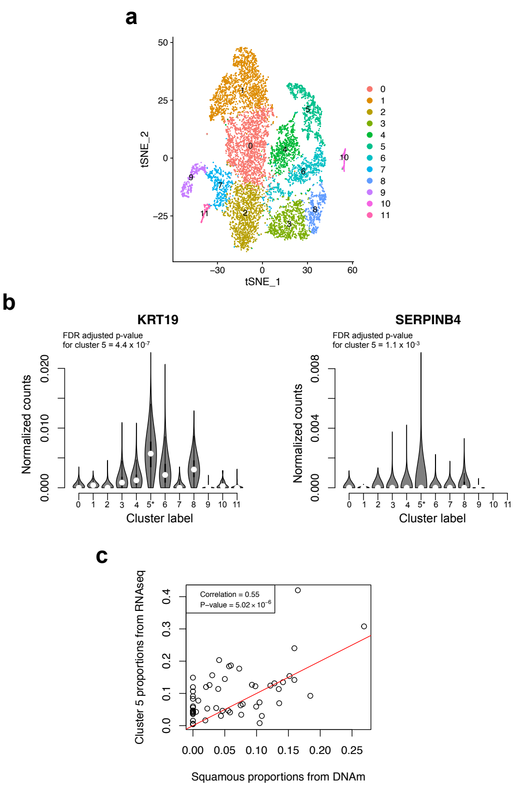


Figure S1: Evidence that "squamous" cell proportion estimates are actually estimates for suprabasal proportions. (**a**): Single cell clusters derived from healthy control nasal brushes in Ziegler et al. [3]. (**b**): Suprabasal marker gene distributions in the aforementioned clusters. P-values were computed using a Wilcoxonrank test. Small p-values indicate the gene is a marker for cluster 5. (**c**): The concordance between DNAm-derived squamous cell proportions and RNAseq-derived cluster 5 proportions.

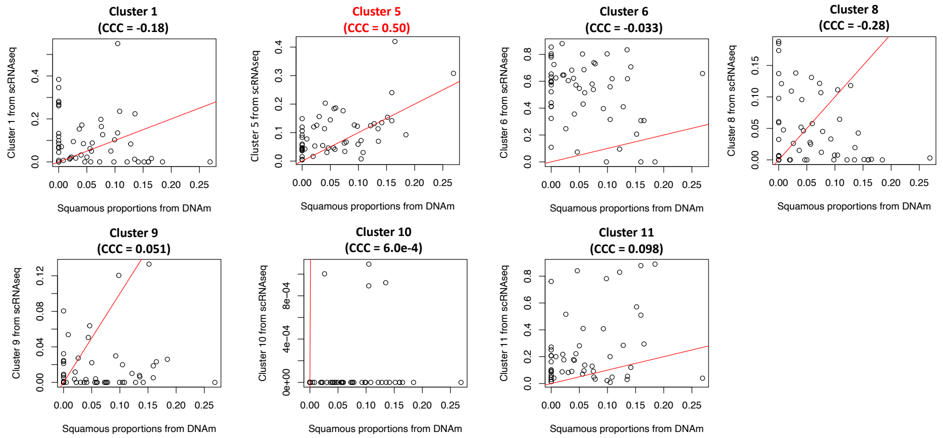


Figure S2: Plots of DNAm-derived "squamous" proportions (x-axes) versus scRNAseq-derived cell type proportions (y-axes). Red lines are the lines $y = x$ and Lin's concordance correlation coefficients (CCC) are indicated. Cluster 5 is the suprabasal cluster. Cell type clusters with proportions equal to zero are not plotted.

| Mean-centered **ciliated** signature | Mean-centered **squamous** signature | Mean-centered **other** signature |
|---|---|---|
| 0.95 (p-value$<10^{-16}$) | 0.86 (p-value$<10^{-16}$) | 0.84 (p-value$<10^{-16}$) |

Table S3: The correlation between the mean-centered columns of the signature matrix ($S_{*k}^{(c)} = S_{*k} - K^{-1}\sum_{k'=1}^{K} S_{*k'}$ for $k = 1, 2, 3$) and the first two left singular vectors ($U \in \mathbb{R}^{p\times 2}$) of the centered DNAm data matrix. The correlation was defined to be $\sup_{u\in \text{image}(U)} \text{corr}\{u, S_{*k}^{(c)}\}$.

#### S3.2 Verifying the signature matrix

The accuracy of the cell type signature derived from external data and described in "Estimating cellular proportions" is critical to the fidelity of our estimates for cellular composition. We therefore evaluated its accuracy using principal components analysis. Specifically, let $p =$#CpGs, $n$ be the samples size, and $K$ be the number of cell types. Let $Y$ be the $p \times n$ data matrix of observed M-values, $S$ be the $p \times K$ signature matrix, and $\Pi$ be the $n \times K$ matrix of cellular proportions (i.e. $\Pi_{ik}$ is the proportion of cell type $k$ in individual $i$). Then if $S$ is accurate, the canonical model relating bulk and single cell omics [8] implies

$$Y = S\Pi^{\top} + E = (K^{-1}S1_K)1_n^{\top} + (SP_{1_K}^{\perp})\Pi^{\top} + E, \tag{S1}$$

where $E$ is a matrix of mean 0 errors, $1_n \in \mathbb{R}^n$ and $1_K \in \mathbb{R}^K$ are vectors of all ones, and $P_{1_K}^{\perp} \in \mathbb{R}^{K\times K}$ is the orthogonal projection matrix that projects vectors onto the orthogonal complement of $1_K$. The second equality in (S1) follows from the fact that the rows of $\Pi$ sum up to one, meaning

$$\Pi = \Pi(K^{-1}1_K1_K^{\top}) + \Pi P_{1_K}^{\perp} = K^{-1}\underbrace{(\Pi 1_K)}_{=1_n}1_K^{\top} + \Pi P_{1_K}^{\perp} = K^{-1}1_n1_K^{\top} + \Pi P_{1_K}^{\perp}.$$

Applying principal components to $Y$ requires first mean-centering its rows, which is equivalent to multiplying it on the right by $P_{1_n}^{\perp}$. This nullifies the first term to the right of the second equality in (S1), meaning the mean-centered $Y$ can be expressed as

$$YP_{1_n}^{\perp} = (SP_{1_K}^{\perp})(P_{1_n}^{\perp}\Pi)^{\top} + \tilde{E},$$

where $\tilde{E} = EP_{1_n}^{\perp}$ is an error matrix containing mean 0 entries.

This analysis shows that if $S$ is an accurate signature matrix, the first left singular vectors of the mean-centered data matrix $Y$ should correlate with the columns of $SP_{1_K}^{\perp}$, where the $k$-th column of $SP_{1_K}^{\perp}$ is $S_{*k} - K^{-1}\sum_{k'=1}^{K} S_{*k'}$. Table S3 shows that the correlation for all $K = 3$ columns is exceedingly large, indicating $S$ is an accurate signature.

#### S3.3 Inferring RSV-related changes in cell proportions

Figure S3 shows that ciliated and other proportions are lower and higher, respectively, in children infected with RSV before 1 year. We used a linear mixed effects model to compute the p-value for these differences. An individual-specific random effect was used to account clustering of some subjects that have cell type estimates at both ages. Since Deprez et al.

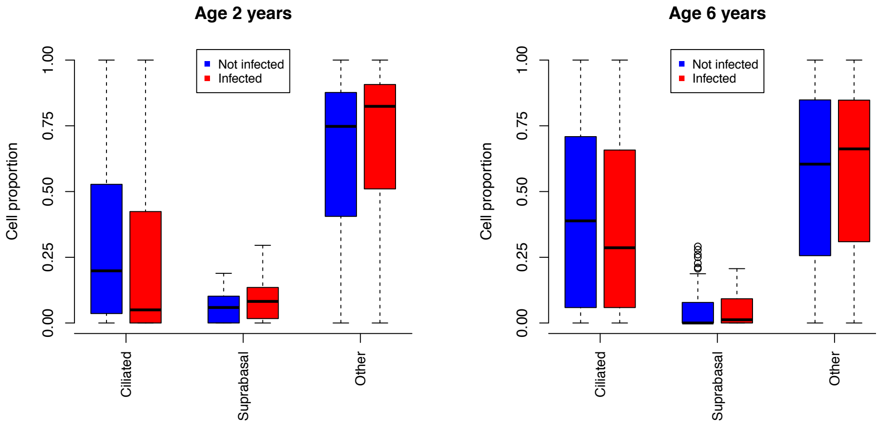


Figure S3: Cell type proportion estimates at each age stratified by whether the subject was infected by RSV by age 1 year.

[2] and the previous subsection suggest other cells are primarily secretory, Persson et al. [9] implies ciliated and other cell proportions should decrease and increase, respectively, upon infection. The aforementioned p-values were therefore one-sided, and were 0.0246 and 0.0410 for the change in ciliated and other proportions, respectively.

#### S3.4 NAEC DNAm is a weighted average of cell type-specific DNAm

As noted in the main text, our cell type-specific analyses rely on the assumption that NAEC DNAm is a weighted average of cell type-specific DNAm, where the weights are cell type proportion. Put another way, the model in (S4) holds. We verified this model holds in Section S3.2.

#### S3.5 The temporal dependence of RSV-related suprabasal DNAm

As alluded to in the main text subsection "Inferring RSV-CpGs in each cell type", Figure S4 shows that the beta coefficients at ages two and six years for the seven RSV-related CpGs identified in suprabasal cells at age two are highly correlated.

The p-value $p^*$ in Figure S4 tests the null hypothesis $H_0$ that suprabasal DNAm at age six at these seven CpGs is independent of RSV infection. To account for the fact that estimates derived from different CpGs may have different error variances, we converted the beta values to z-scores by standardizing them with their standard errors. Let $\hat{z}_{gt}$ be the z-score at CpG $g = 1, \ldots, 7$ and time $t = 1, 2$ (time 1:=2 years, time 2:= 6 years), where we use "$\hat{\cdot}$" to emphasize observed z-scores are random functions of the data. We defined a multivariate test statistic to be $(S, P)$, where $S$ is the number of CpGs (out of 7) whose z-scores at times 1 and 2 had the same sign and $P$ is the p-value from the regression of z-scores at time 2 onto those at time 1. If $H_0$ were true, then we would expect little-to-no concordance between z-scores, meaning large values of $S$ and small values of $P$ provide evidence against $H_0$. We therefore defined $p^*$ to be

$$p^* = \text{pr}(S \geq s, P \leq p \mid \{\hat{z}_{g1}\}_{g=1}^{7}),$$

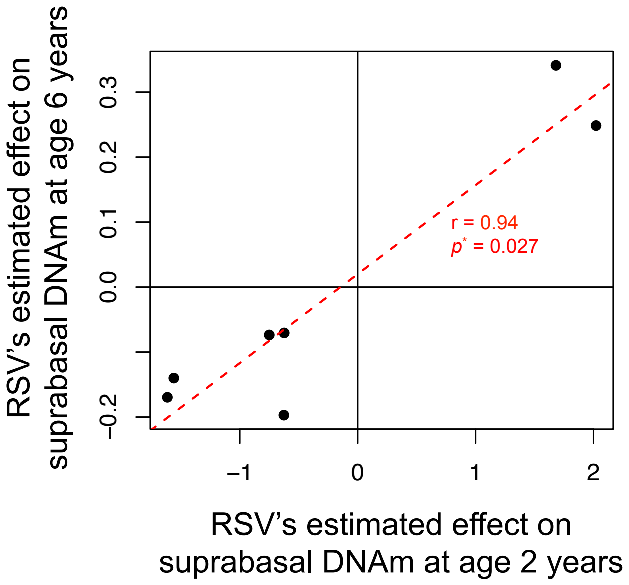


Figure S4: The beta coefficients at ages two and six years for the seven RSV-related CpGs identified in suprabasal cells at age two. The dashed red line is the line of best fit.

where $s = 7$ and $p = 8.7\text{e-}4$ were the values observed in our dataset. We conditioned on the z-scores observed at time 1 because our selection of the above seven CpGs was based those z-scores.

Computing $p^*$ is straightforward if z-scores at different times are independent. However, since 48 children had DNAm samples at both ages, z-scores may be temporally dependent. To adjust for this dependence, we first note that conditional on $\{\hat{z}_{g1}\}_{g=1}^7$, $S$ and $P$ are functions of $\{\hat{z}_{g2}\}_{g=1}^7$. Therefore, it suffices to determine $\text{pr}(\{\hat{z}_{g2}\}_{g=1}^7 \mid \{\hat{z}_{g1}\}_{g=1}^7)$ under the null hypothesis $H_0$ and derive $p^*$ by Monte Carlo. To do so, we first note that because our regressions adjust for latent confounds (see Methods), McKennan et al. [10] implies it is appropriate to assume that z-scores for different CpGs are independent, i.e.

$$\text{pr}(\{\hat{z}_{g2}\}_{g=1}^7 \mid \{\hat{z}_{g1}\}_{g=1}^7) = \prod_{g=1}^{p} \text{pr}(\hat{z}_{g2} \mid \hat{z}_{g1}).$$

Next, since $\hat{z}_{gt}$ are z-scores, they are normally distributed with unit variance:

$$(\hat{z}_{g1}, \hat{z}_{g2})^\top \sim N\left((\mu_{g1}, \mu_{g2})^\top, \begin{pmatrix} 1 & \rho_g \\ \rho_g & 1 \end{pmatrix}\right), \quad g = 1, \ldots, 7,$$

where we estimated $\rho_g = \rho = 0.37$ in our data. Since $\mu_{gt} = 0$ if RSV infection has no effect on CpG $g$'s suprabasal DNAm at time $t$, $\mu_{g2} = 0$ for all $g$ under the null hypothesis $H_0$. Since $\mu_{g1}$ is unknown and may not be zero under $H_0$, we follow Stephens [11] and McKennan et al. [12] and model it as a mixture of normal distributions:

$$\mu_{g1} \sim \textstyle\sum_k \pi_k N(0, \sigma_k^2), \quad \pi_k \in [0,1], \quad \textstyle\sum_k \pi_k = 1, \quad \sigma_k^2 \geq 0, \quad g = 1, \ldots, 7,$$

where we used all CpGs and the empirical Bayes algorithm in Stephens [11] to estimate the hyperparameters $\pi_k$ and $\sigma_k^2$. Assuming the null hypothesis $H_0$ was true, $\hat{z}_{g2}$ can therefore be sampled conditional on $\hat{z}_{g1}$ as follows:

$$e_{g1} \mid \hat{z}_{g1} \sim \textstyle\sum_k \tilde{\pi}_k(\hat{z}_{g1}) N(\hat{z}_{g1}/(\sigma_k^2+1), \sigma_k^2/(\sigma_k^2+1)), \quad \tilde{\pi}_k(\hat{z}_{g1}) = \frac{\pi_k \mathcal{N}(\hat{z}_{g1};0,\sigma_k^2+1)}{\sum_{k'} \pi_{k'} \mathcal{N}(\hat{z}_{g1};0,\sigma_{k'}^2+1)}$$
$$\hat{z}_{g2} \mid e_{g1} \sim N(\rho e_{g1}, 1-\rho^2).$$

We used this procedure to simulate $10^4$ datasets $\{\hat{z}_{g2}\}_{g=1}^7$ under $H_0$, which we then use in our Monte Carlo to compute $p^*$.

#### S3.6 Cell type-specific correlation between DNAm and gene expression

Our methodology is based on existing models relating bulk and cell type-specific 'omic' data [8, 13]. Fix a CpG, gene pair and let $D_i^{(B)}$ ($G_i^{(B)}$) and $D_{ki}^{(CTS)}$ ($G_{ki}^{(CTS)}$) be subject $i$'s observed bulk DNAm (gene expression) and unobserved cell type-specific DNAm (gene expression) in cell type $k$, and let $\pi_{ki}$ be cell type $k$'s proportion in subject $i$, which was estimated in Section S3.1. Our goal is to estimate the cell type-specific covariance between DNAm and gene expression, i.e. $\rho_k = \text{Cov}\{D_{ki}^{(CTS)}, G_{ki}^{(CTS)}\}$ for each cell type $k$. First, $D_i^{(B)} = \sum_k \pi_{ki} D_{ki}^{(CTS)} + \epsilon_{D,i}$ and $G_i^{(B)} = \sum_k \pi_{ki} G_{ki}^{(CTS)} + \epsilon_{G,i}$ where $\epsilon_{D,i}$ and $\epsilon_{G,i}$ are independent mean-zero error terms [8]. Then assuming omics from different cell types are independent [13],

$$\mathbb{E}\{R_{D,i}^{(B)} R_{G,i}^{(B)}\} = \text{Cov}\{D_i^{(B)}, G_i^{(B)}\} = \textstyle\sum_k \pi_{ki}^2 \,\text{Cov}\{D_{ki}^{(CTS)}, G_{ki}^{(CTS)}\} = \sum_k \pi_{ki}^2 \rho_k \qquad \text{(S2)}$$

where $R_{D,i}^{(B)}$ and $R_{G,i}^{(B)}$ are the residuals formed by mean-centering $D_i^{(B)}$ and $G_i^{(B)}$. We derived the residuals by regressing out the observed covariates and latent factors discussed in Methods from $D_i^{(B)}$ and $G_i^{(B)}$.

We used the method of moments estimator from Kerin et al. [14] to estimate $\rho_k$. To describe the estimator, let $Y = (R_{D,1}^{(B)} R_{G,1}^{(B)}, \ldots, R_{D,n}^{(B)} R_{G,n}^{(B)})^\top \in \mathbb{R}^n$, $\rho = (\rho_1, \ldots, \rho_K)^\top \in \mathbb{R}^K$, and $\Pi^\top = (\pi_{ki}^2) \in \mathbb{R}^{K \times n}$ be the matrix of squared cellular proportions. Then (S2) implies

$$Y = \Pi\rho + \delta, \quad \mathbb{E}(\delta) = 0,$$

meaning the ordinary least squares estimator, $\hat{\rho} = (\Pi^\top\Pi)^{-1}\Pi^\top Y$, is an unbiased estimator for $\rho$. While the entries of $Y$ are independent, they may not have the same variance, meaning we must use the sandwich variance estimator [15] to estimate $\text{Var}(\hat{\rho})$. This can be expressed as

$$\hat{V}(\hat{\rho}) = (\Pi^\top\Pi)^{-1} \left\{ \sum_{i=1}^n (Y_i - \Pi_{i*}^\top \hat{\rho})^2 \Pi_{i*} \Pi_{i*}^\top \right\} (\Pi^\top\Pi)^{-1},$$

where $\Pi_{i*} \in \mathbb{R}^K$ is the $i$-th row of $\Pi$. We subsequently use the normal approximation to compute p-values for the entries for $\rho$.

#### S3.7 The dependence between suprabasal DNAm and childhood wheeze

We used the approach taken in Zheng et al. [16] to determine the association between suprabasal gene expression and wheeze at age four years, which is analogous to regressing cell type-specific (CTS) gene expression onto wheeze (yes/no). This is the reverse of the regression we would ideally perform if we believed CTS expression had an effect on wheeze risk. We take this approach for two reasons. The first is because the methodology to regress a binary variable (wheeze) onto CTS expression does not exist. Notably, the method proposed in Rahmani et al. [17], which to our knowledge is the only work to consider regressing a dependent variable onto unobserved CTS expression, requires the dependent variable be normally distributed. Extending their work to binary dependent variables is non-trivial, as they rely on the normality assumption to derive conditional distributions. The second reason is because the reverse regression allows us to account for latent confounders that may confound the relationship between CTS expression and wheeze risk [18]. The statistical guarantees from McKennan et al. [18] only hold in the reverse setting.

We next provide a mathematical justification for our approach. Our goal is to show that we can use the reverse regression to infer whether the expression in a particular cell type increases, decreases, or has no impact on wheeze risk. To do so, let $\boldsymbol{D}_i^{(\text{CTS})} \in \mathbb{R}^K$ be subject $i$'s unobserved vector of CTS expression in $K$ cell types at a CpG. We also let $\boldsymbol{x}_i$ be a vector of nuisance covariates, which include cell type proportions and other factors we would like to adjust for. We assume wheeze ($W_i$) follows a generalized linear model, where wheeze risk is dependent on CTS expression and the nuisance covariates:

$$\mathbb{P}\{W_i = 1 \mid \boldsymbol{D}_i^{(\text{CTS})}, \boldsymbol{x}_i\} = g\{\mu_W + \boldsymbol{\beta}^\top \boldsymbol{D}_i^{(\text{CTS})} + \boldsymbol{\gamma}^\top \boldsymbol{x}_i\}, \tag{S3}$$

where the function $g(\cdot)$ is the inverse of the link function. For example, $g$ would be the "expit" function in logistic regression. The elements of $\boldsymbol{\beta} \in \mathbb{R}^K$ dictate the impact of CTS expression on wheeze risk, where expression in the $k$-th cell type increases (decreases) wheeze risk if $\boldsymbol{\beta}_k > 0$ ($\boldsymbol{\beta}_k < 0$). To explore what happens when we reverse the regression, we follow Cai et al. [13] and Rahmani et al. [17] and model CTS expression as

$$\boldsymbol{D}_i^{(\text{CTS})} \mid \boldsymbol{x}_i \sim N_K(\boldsymbol{\mu}_D + \boldsymbol{\Gamma}\boldsymbol{x}_i, \boldsymbol{\Sigma}), \quad \boldsymbol{\Sigma} = \text{diagonal } K \times K \text{ matrix}, \tag{S4}$$

where $\boldsymbol{\mu}_D \in \mathbb{R}^K$ and the matrix of coefficients $\boldsymbol{\Gamma}$ dictates the relationship between CTS expression and the nuisance covariates $\boldsymbol{x}_i$. Note we by defining $\tilde{\boldsymbol{D}}_i^{(\text{CTS})} = \boldsymbol{D}_i^{(\text{CTS})} - \mathbb{E}\{\boldsymbol{D}_i^{(\text{CTS})} \mid \boldsymbol{x}_i\}$, we can re-write (S3) as

$$\begin{aligned}&\mathbb{P}\{W_i = 1 \mid \boldsymbol{D}_i^{(\text{CTS})}, \boldsymbol{x}_i\} = g\{\tilde{\mu}_W + \boldsymbol{\beta}^\top \tilde{\boldsymbol{D}}_i^{(\text{CTS})} + \tilde{\boldsymbol{\gamma}}^\top \boldsymbol{x}_i\}\\ &\tilde{\boldsymbol{D}}_i^{(\text{CTS})} \mid \boldsymbol{x}_i \sim N_K(0, \boldsymbol{\Sigma}), \quad \tilde{\mu}_W = \mu_W + \boldsymbol{\beta}^\top \boldsymbol{\mu}_D, \quad \tilde{\boldsymbol{\gamma}} = \boldsymbol{\gamma} + \boldsymbol{\Gamma}^\top \boldsymbol{\beta}.\end{aligned}$$

A justification for reversing the regression requires showing the following:

1) If $\boldsymbol{\beta}_k = 0$, then $\mathbb{E}\{\boldsymbol{D}_{ik}^{(\text{CTS})} \mid W_i, \boldsymbol{x}_i\} = \mathbb{E}\{\boldsymbol{D}_{ik}^{(\text{CTS})} \mid \boldsymbol{x}_i\}$ does not depend on $W_i$ and is a linear function of $\boldsymbol{x}_i$.

2) $\text{Var}\{\boldsymbol{D}_i^{(\text{CTS})} \mid W_i, \boldsymbol{x}_i\}$ does not depend on $i$.

3) If $\boldsymbol{\beta}_k > 0$, then $\mathbb{E}\{\boldsymbol{D}_{ik}^{(\text{CTS})} \mid W_i, \boldsymbol{x}_i\}$ is a linear function of $W_i$ and $\boldsymbol{x}_i$ and $\mathbb{E}\{\boldsymbol{D}_{ik}^{(\text{CTS})} \mid W_i = 1, \boldsymbol{x}_i\} > \mathbb{E}\{\boldsymbol{D}_{ik}^{(\text{CTS})} \mid W_i = 0, \boldsymbol{x}_i\}$. This will allow us to infer the direction of the effect.

Showing 1), 2), and 3) will allow us to use Zheng et al. [16] to test whether expression at cell type $k$ impacts wheeze risk and, if it does, also infer the direction of association. To show 1), suppose $\boldsymbol{\beta}_k = 0$. Then (S4) implies

$$\mathbb{E}\{\boldsymbol{D}_{ik}^{(\text{CTS})} \mid W_i, \boldsymbol{x}_i\} = \mathbb{E}\{\tilde{\boldsymbol{D}}_{ik}^{(\text{CTS})} \mid W_i, \boldsymbol{x}_i\} + \boldsymbol{\mu}_{Dk} + \boldsymbol{\Gamma}_{k*}^\top \boldsymbol{x}_i = 0 + \boldsymbol{\mu}_{Dk} + \boldsymbol{\Gamma}_{k*}^\top \boldsymbol{x}_i,$$

where $\boldsymbol{\mu}_{Dk}$ and $\boldsymbol{\Gamma}_{k*}$ are the $k$-th element and row of $\boldsymbol{\mu}_D$ and $\boldsymbol{\Gamma}$, respectively. The second equality follows from the fact that the expression in cell type $k$ is independent of $W_i$ if $\boldsymbol{\beta}_k = 0$.

To show 2), we assume the elements of $\boldsymbol{\beta}$ in (S3) are zero or relatively small, i.e. the impact of CTS expression on wheeze risk is relatively small. This is a common assumption used to develop statistical methods designed to study asthma- and allergy-related traits [18, 10, 1]. We can then use a Taylor expansion to show 2). In the below derivation, we let $\eta_i = \tilde{\mu}_W + \tilde{\boldsymbol{\gamma}}^\top \boldsymbol{x}_i$.

$$\begin{aligned}
\text{Var}\{\boldsymbol{D}_i^{(\text{CTS})} \mid W_i, \boldsymbol{x}_i\} &= \text{Var}\{\tilde{\boldsymbol{D}}_i^{(\text{CTS})} \mid W_i, \boldsymbol{x}_i\} \\
&= \frac{\int \tilde{\boldsymbol{D}}_i^{(\text{CTS})}\{\tilde{\boldsymbol{D}}_i^{(\text{CTS})}\}^\top g\{\eta_i + \boldsymbol{\beta}^\top \tilde{\boldsymbol{D}}_i^{(\text{CTS})}\} \,\text{pr}\{\tilde{\boldsymbol{D}}_i^{(\text{CTS})}\} d\tilde{\boldsymbol{D}}_i^{(\text{CTS})}}{\int g\{\eta_i + \boldsymbol{\beta}^\top \tilde{\boldsymbol{D}}_i^{(\text{CTS})}\} \,\text{pr}\{\tilde{\boldsymbol{D}}_i^{(\text{CTS})}\} d\tilde{\boldsymbol{D}}_i^{(\text{CTS})}} \\
&= \frac{g(\eta_i) \int \tilde{\boldsymbol{D}}_i^{(\text{CTS})}\{\tilde{\boldsymbol{D}}_i^{(\text{CTS})}\}^\top \,\text{pr}\{\tilde{\boldsymbol{D}}_i^{(\text{CTS})}\} d\tilde{\boldsymbol{D}}_i^{(\text{CTS})} + O(\|\boldsymbol{\beta}\|^2)}{g(\eta_i) + O(\|\boldsymbol{\beta}\|^2)} \\
&= \frac{g(\eta_i)\boldsymbol{\Sigma} + O(\|\boldsymbol{\beta}\|^2)}{g(\eta_i) + O(\|\boldsymbol{\beta}\|^2)} = \boldsymbol{\Sigma} + O(\|\boldsymbol{\beta}\|^2),
\end{aligned}$$

which is approximately equal to $\boldsymbol{\Sigma}$ when the entries of $\boldsymbol{\beta}$ are small, and therefore shows 2).

To show 3) we make an additional assumption requiring the elements of $\boldsymbol{\gamma}$ in (S3) are moderate to small. That is, the impact of nuisance covariates on wheeze risk is not large and a moderate to large sample size is required to be able to infer their effects. This is consistent with existing epidemiological studies, which often require hundreds to thousands of samples to infer risk factors for wheeze and other related phenotypes [19, 20, 21]. We first note that

$$\mathbb{E}\{\boldsymbol{D}_i^{(\text{CTS})} \mid W_i, \boldsymbol{x}_i\} = \mathbb{E}\{\tilde{\boldsymbol{D}}_i^{(\text{CTS})} \mid W_i, \boldsymbol{x}_i\} + \boldsymbol{\mu}_D + \boldsymbol{\Gamma}\boldsymbol{x}_i,$$

meaning it suffices to show that $\mathbb{E}\{\tilde{\boldsymbol{D}}_i^{(\text{CTS})} \mid W_i, \boldsymbol{x}_i\}$ is a linear function of $W_i$. Let $s_i = 2W_i - 1$ and again define $\eta_i = \tilde{\mu}_W + \tilde{\boldsymbol{\gamma}}_i^\top \boldsymbol{x}$. Then

$$\begin{aligned}
\mathbb{E}\{\tilde{\boldsymbol{D}}_i^{(\text{CTS})} \mid W_i, \boldsymbol{x}_i\} &= s_i \frac{\int \tilde{\boldsymbol{D}}_i^{(\text{CTS})} g\{\eta_i + \boldsymbol{\beta}^\top \tilde{\boldsymbol{D}}_i^{(\text{CTS})}\} \,\text{pr}\{\tilde{\boldsymbol{D}}_i^{(\text{CTS})}\} d\tilde{\boldsymbol{D}}_i^{(\text{CTS})}}{\int g\{\eta_i + \boldsymbol{\beta}^\top \tilde{\boldsymbol{D}}_i^{(\text{CTS})}\} \,\text{pr}\{\tilde{\boldsymbol{D}}_i^{(\text{CTS})}\} d\tilde{\boldsymbol{D}}_i^{(\text{CTS})}} \\
&= s_i \frac{\dot{g}(\eta_i) \int \tilde{\boldsymbol{D}}_i^{(\text{CTS})}\{\tilde{\boldsymbol{D}}_i^{(\text{CTS})}\}^\top \boldsymbol{\beta} \,\text{pr}\{\tilde{\boldsymbol{D}}_i^{(\text{CTS})}\} d\tilde{\boldsymbol{D}}_i^{(\text{CTS})} + O(\|\boldsymbol{\beta}\|^3)}{g(\mu_i) + O(\|\boldsymbol{\beta}\|^2)}
\end{aligned}$$

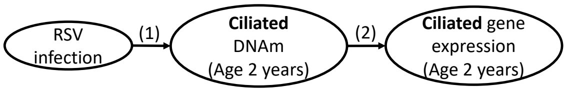


| CpG | Nearest gene (CpG location) | Direction of arrow (1)'s effect | Direction of arrow (2)'s effect | Indirect effect direction | Corroborating evidence |
|---|---|---|---|---|---|
| cg23865736 | SBNO2 (Gene body) | ↑ | ↓ | ↓ | SBNO2 is a TF that suppresses the transcription of NF-κB and other inflammatory genes.[1] Its DNAm in NAECs is altered upon RSV immunoprophylaxis.[2] |
| cg22997144 | SEPT9 (Other) | ↑ | ↓ | ↓ | ↓SEPT9 and RSV infection have both been shown to impair epithelial wound healing.[3,4] |
| cg18114394 | DTX1 (Other) | ↑ | ↓ | ↓ | DTX1 promotes T cell anergy and reduces T cell activation.[5] |
| cg22510620 | PGAP3 (Gene body) | ↑ | ↓ | ↓ | GWAS and eQTL evidence suggest PGAP3 is elevated in the epithelial cells of patients with inflammatory diseases.[6] |
| cg22084244 | ARRB2 (TSS) | ↑ | ↑ | ↑ | Promotes Th2 chemotaxis.[7] |

Table S4: Results and functional interpretation for ciliated RSV-CpGs at age 2 years whose DNAm was correlated with its nearest gene's expression in suprabasal cells at an FDR of 25%. The indirect effect is the product of the directions of arrows (1) and (2). [1]: El Kasmi et al. [24]; [2]: Xu et al. [25]; [3]: Ivanov et al. [26]; [4]: Andersson et al. [27]; [5]: Hsiao et al. [28]; [6]: Yuan et al. [23]; [7]: Lin et al. [29].

$$
\begin{aligned}
&= s_i \frac{\dot{g}(\eta_i)}{g(\eta_i)} \boldsymbol{\Sigma}\boldsymbol{\beta} + O(\|\boldsymbol{\beta}\|^3) \\
&= (2W_i - 1) \frac{\dot{g}(\tilde{\mu}_W)}{g(\tilde{\mu}_W)} \boldsymbol{\Sigma}\boldsymbol{\beta} + O\{\|\boldsymbol{\beta}\|(\|\boldsymbol{\beta}\|^2 + \|\boldsymbol{\gamma}\|)\},
\end{aligned}
$$

which is a linear function of $W_i$. What's more, since $\boldsymbol{\Sigma}$ is diagonal and $\frac{2\dot{g}(\tilde{\mu}_W)}{g(\tilde{\mu}_W)} > 0$,

$$
\mathbb{E}\{\boldsymbol{D}_{ik}^{(\mathrm{CTS})} \mid W_i = 1, \boldsymbol{x}_i\} > \mathbb{E}\{\boldsymbol{D}_{ik}^{(\mathrm{CTS})} \mid W_i = 0, \boldsymbol{x}_i\} \text{ if } \boldsymbol{\beta}_k > 0.
$$

This shows 3).

### S4 Supplemental results

##### The correlation between DNAm and gene expression in ciliated cells

Table S4 reports the ciliated cell-specific correlation between the DNAm at ciliated RSV-CpGs and the expression of neighboring genes, along with a existing work that corroborates the implied indirect effect of RSV infection on ciliated gene expression. Perhaps the most surprising result is that of PGAP3, which suggests that DNAm mediates an RSV-induced reduction in PGAP3 levels in ciliated cells. While the direction of this relationship is the opposite of what we might expect given the expression quantitative trait loci (eQTL) and childhood asthma genome-wide association study (GWAS) evidence for single nucleotide polymorphisms (SNPs) in PGAP3 [22], evidence from other inflammatory diseases, including ulcerative colitis and inflammatory bowel disease [23], are congruent with out results (Tab. S4).

### References


[1] A. Morin, E. E. Thompson, B. A. Helling, et al. “A functional genomics pipeline to identify high-value asthma and allergy CpGs in the human methylome”. In: *Journal of Allergy and Clinical Immunology* 151.6 (June 2023), pp. 1609–1621. DOI: 10.1016/j.jaci.2022.12.828.

[2] M. Deprez, L.-E. Zaragosi, M. Truchi, et al. “A Single-Cell Atlas of the Human Healthy Airways”. In: *American Journal of Respiratory and Critical Care Medicine* 202.12 (Dec. 2020), pp. 1636–1645. DOI: 10.1164/rccm.201911-2199oc. URL: https://doi.org/10.1164/rccm.201911-2199oc.

[3] C. G. K. Ziegler, V. N. Miao, A. H. Owings, et al. “Impaired local intrinsic immunity to SARS-CoV-2 infection in severe COVID-19”. In: *Cell* 184.18 (2021), 4713–4733.e22.

[4] M. Karlsson, C. Zhang, L. Méar, et al. “A single–cell type transcriptomics map of human tissues”. In: *Science Advances* 7.31 (July 2021). ISSN: 2375-2548. DOI: 10.1126/sciadv.abh2169. URL: http://dx.doi.org/10.1126/sciadv.abh2169.

[5] X. Wang, J. Park, K. Susztak, et al. “Bulk tissue cell type deconvolution with multi-subject single-cell expression reference”. In: *Nature Communications* 10.1 (2019), p. 380. DOI: 10.1038/s41467-018-08023-x.

[6] L. I.-K. Lin. “A Concordance Correlation Coefficient to Evaluate Reproducibility”. In: *Biometrics* 45.1 (Mar. 1989), p. 255. ISSN: 0006-341X. DOI: 10.2307/2532051. URL: http://dx.doi.org/10.2307/2532051.

[7] F. A. Vieira Braga, G. Kar, M. Berg, et al. “A cellular census of human lungs identifies novel cell states in health and in asthma”. In: *Nature Medicine* 25.7 (2019), pp. 1153–1163.

[8] M. Cai, M. Yue, T. Chen, et al. “Robust and accurate estimation of cellular fraction from tissue omics data via ensemble deconvolution”. In: *Bioinformatics* 38.11 (Apr. 2022). Ed. by I. Birol, pp. 3004–3010. DOI: 10.1093/bioinformatics/btac279.

[9] B. D. Persson, A. B. Jaffe, R. Fearns, et al. “Respiratory Syncytial Virus Can Infect Basal Cells and Alter Human Airway Epithelial Differentiation”. In: *PLoS ONE* 9.7 (July 2014). Ed. by K. Harrod, e102368. DOI: 10.1371/journal.pone.0102368. URL: https://doi.org/10.1371/journal.pone.0102368.

[10] C. McKennan and D. Nicolae. “Estimating and Accounting for Unobserved Covariates in High-Dimensional Correlated Data”. In: *Journal of the American Statistical Association* 117.537 (June 2020), pp. 225–236. DOI: 10.1080/01621459.2020.1769635. URL: https://doi.org/10.1080/01621459.2020.1769635.

[11] M. Stephens. “False discovery rates: a new deal”. In: *Biostatistics* (Oct. 2016), kxw041. DOI: 10.1093/biostatistics/kxw041.

[12] C. McKennan, K. Naughton, C. Stanhope, et al. “Longitudinal data reveal strong genetic and weak non-genetic components of ethnicity-dependent blood DNA methylation levels”. In: *Epigenetics* 16.6 (Sept. 2020), pp. 662–676. DOI: 10.1080/15592294.2020.1817290. URL: https://doi.org/10.1080/15592294.2020.1817290.

[13] B. Cai, J. Zhang, H. Li, et al. *Statistical Inference of Cell-type Proportions Estimated from Bulk Expression Data*. 2022. arXiv: 2209.04038 [stat.ME].

[14] M. Kerin and J. Marchini. “Inferring Gene-by-Environment Interactions with a Bayesian Whole-Genome Regression Model”. In: *The American Journal of Human Genetics* 107.4 (Oct. 2020), pp. 698–713. DOI: 10.1016/j.ajhg.2020.08.009. URL: https://doi.org/10.1016/j.ajhg.2020.08.009.

[15] G. Kauermann and R. J. Carroll. “A Note on the Efficiency of Sandwich Covariance Matrix Estimation”. In: *Journal of the American Statistical Association* 96.456 (Dec. 2001), pp. 1387–1396. DOI: 10.1198/016214501753382309. URL: https://doi.org/10.1198/016214501753382309.

[16] S. C. Zheng, C. E. Breeze, S. Beck, et al. “Identification of differentially methylated cell types in epigenome-wide association studies”. In: *Nature Methods* 15.12 (2018), pp. 1059–1066.

[17] E. Rahmani, R. Schweiger, B. Rhead, et al. “Cell-type-specific resolution epigenetics without the need for cell sorting or single-cell biology”. In: *Nature Communications* 10.1 (July 2019). DOI: 10.1038/s41467-019-11052-9. URL: https://doi.org/10.1038/s41467-019-11052-9.

[18] C. McKennan and D. Nicolae. “Accounting for unobserved covariates with varying degrees of estimability in high-dimensional biological data”. In: *Biometrika* 106.4 (Sept. 2019), pp. 823–840. ISSN: 0006-3444. DOI: 10.1093/biomet/asz037.

[19] D. J. Jackson, R. E. Gangnon, M. D. Evans, et al. “Wheezing Rhinovirus Illnesses in Early Life Predict Asthma Development in High-Risk Children”. In: *American Journal of Respiratory and Critical Care Medicine* 178.7 (Oct. 2008), pp. 667–672. DOI: 10.1164/rccm.200802-309oc. URL: https://doi.org/10.1164/rccm.200802-309oc.

[20] P. Wu, W. D. Dupont, M. R. Griffin, et al. “Evidence of a Causal Role of Winter Virus Infection during Infancy in Early Childhood Asthma”. In: *American Journal of Respiratory and Critical Care Medicine* 178.11 (Dec. 2008), pp. 1123–1129. DOI: 10.1164/rccm.200804-579oc. URL: https://doi.org/10.1164/rccm.200804-579oc.

[21] C. Rosas-Salazar, T. Chirkova, T. Gebretsadik, et al. “Respiratory syncytial virus infection during infancy and asthma during childhood in the USA (INSPIRE): a population-based, prospective birth cohort study”. In: *The Lancet* 401.10389 (May 2023), pp. 1669–1680. DOI: 10.1016/s0140-6736(23)00811-5. URL: https://doi.org/10.1016/s0140-6736(23)00811-5.

[22] C. Ober, C. G. McKennan, K. M. Magnaye, et al. “Expression quantitative trait locus fine mapping of the 17q12–21 asthma locus in African American children: a genetic association and gene expression study”. In: *The Lancet Respiratory Medicine* 8.5 (May 2020), pp. 482–492. DOI: 10.1016/s2213-2600(20)30011-4. URL: https://doi.org/10.1016/s2213-2600(20)30011-4.

[23] F. Yuan, R. J. Hung, N. Walsh, et al. "Genome-Wide Association Study Data Reveal Genetic Susceptibility to Chronic Inflammatory Intestinal Diseases and Pancreatic Ductal Adenocarcinoma Risk". In: *Cancer Research* 80.18 (Sept. 2020), pp. 4004–4013. DOI: 10.1158/0008-5472.can-20-0447. URL: https://doi.org/10.1158/0008-5472.can-20-0447.

[24] K. C. El Kasmi, A. M. Smith, L. Williams, et al. "Cutting Edge: A Transcriptional Repressor and Corepressor Induced by the STAT3-Regulated Anti-Inflammatory Signaling Pathway1". In: *The Journal of Immunology* 179.11 (Dec. 2007), pp. 7215–7219. ISSN: 0022-1767. DOI: 10.4049/jimmunol.179.11.7215. URL: https://doi.org/10.4049/jimmunol.179.11.7215.

[25] C.-J. Xu, N. M. Scheltema, C. Qi, et al. "Infant RSV immunoprophylaxis changes nasal epithelial DNA methylation at 6 years of age". In: *Pediatric Pulmonology* 56.12 (Sept. 2021), pp. 3822–3831. DOI: 10.1002/ppul.25643. URL: https://doi.org/10.1002/ppul.25643.

[26] A. I. Ivanov, H. T. Le, N. G. Naydenov, et al. "Novel Functions of the Septin Cytoskeleton". In: *The American Journal of Pathology* 191.1 (Jan. 2021), pp. 40–51. DOI: 10.1016/j.ajpath.2020.09.007. URL: https://doi.org/10.1016/j.ajpath.2020.09.007.

[27] C. K. Andersson, J. Iwasaki, J. Cook, et al. "Impaired airway epithelial cell wound-healing capacity is associated with airway remodelling following RSV infection in severe preschool wheeze". In: *Allergy* 75.12 (July 2020), pp. 3195–3207. DOI: 10.1111/all.14466. URL: https://doi.org/10.1111/all.14466.

[28] H.-W. Hsiao, W.-H. Liu, C.-J. Wang, et al. "Deltex1 Is a Target of the Transcription Factor NFAT that Promotes T Cell Anergy". In: *Immunity* 31.1 (July 2009), pp. 72–83. DOI: 10.1016/j.immuni.2009.04.017. URL: https://doi.org/10.1016/j.immuni.2009.04.017.

[29] R. Lin, Y. ho Choi, D. A. Zidar, et al. "$\beta$-Arrestin-2–Dependent Signaling Promotes CCR4–mediated Chemotaxis of Murine T-Helper Type 2 Cells". In: *American Journal of Respiratory Cell and Molecular Biology* 58.6 (June 2018), pp. 745–755. DOI: 10.1165/rcmb.2017-0240oc. URL: https://doi.org/10.1165/rcmb.2017-0240oc.
